# Translation-specific disruption of Col1a1 expression in multiple models of Spinal Muscular Atrophy can be rescued by Risdiplam

**DOI:** 10.1101/2025.06.03.657600

**Authors:** Gaurav Sharma, Martina Paganin, Yu-Ting Huang, Elena Parenthaler, Ilaria Signoria, Kiterie M.E. Faller, Federica Maniscalco, Cecilia Perrucci, Deborah Donzel, Helena Chaytow, Manuela Basso, Fabio Lauria, Rashmi Kothary, W Ludo van der Pol, Peter Claus, Ewout Groen, Thomas H. Gillingwater, Gabriella Viero

## Abstract

Spinal muscular atrophy (SMA) is a monogenic neurodegenerative disorder caused by decreased levels of Survival of Motor Neuron (SMN) protein. If left untreated, SMA patients have a poor prognosis, marked by the degeneration of motor neurons, progressive muscle weakness and atrophy. The approval of SMN-restoring therapies that improve symptoms and lifespan in patients with SMA has created emerging, non-neuronal phenotypes and an urgent need for deepening our understanding of disease pathogenesis.

Leveraging the knowledge that SMN loss drives alterations in translation, we used multiple tissues from a mouse model of SMA to uncover early translational alterations in key mRNAs and proteins, which act as contributors to pathogenesis and hallmarks of the disease. Among hundreds of differentially translated mRNAs, *Col1a1* emerged as a translation-specific manifestation of early defects in the mouse model. These findings were confirmed in fibroblasts derived from patients with varying levels of disease severity. Notably, treatment with SMN-restoring therapies rescued COL1A1 protein levels, particularly in fibroblasts from patients with the most severe forms of the disease.

Overall, our study identifies COL1A1 as an indicator of disease severity in SMA, which captures early molecular alterations and respond to SMN-modifying therapies.

## Introduction

Spinal Muscular Atrophy (SMA) is a monogenic disease caused by deletions or mutations in the *SMN1* gene (*Lefebvre et al., 1995*), which lead to low levels of SMN protein (*Lefebvre et al., 1997*). Humans have multiple copies of an *SMN1* gene paralogue, namely *SMN2.* This gene presents with single nucleotide polymorphisms, of which a C-to-T transition in exon 7 favours exon 7 skipping in about 90% of transcripts (*Monani et al., 1999*). This event results in the majority of *SMN2* mRNA being translated into a truncated protein that is rapidly degraded (*Vitte et al., 2007*). Only a minority of *SMN2* mRNA is correctly spliced and translated into a functional SMN protein, making *SMN2* unable to fully compensate for the loss of *SMN1* in SMA patients (*Lefebre et al., 1997; Lorson et al., 1999*). Leveraging the potential to correct *SMN2* splicing or to exogenously compensate the *SMN1* gene loss through gene therapy, three disease-modifying therapies (DMTs) that restore SMN protein levels have been successfully introduced in clinical practice (*Finkel et al., 2017; Mendell et al., 2017; Poirer et al., 2018*). These therapies, if delivered in a timely manner, effectively alter disease progression and prolong patients’ survival but do not cure SMA (*Nishio et al., 2023; Chaytow et al., 2021*). Importantly, DMTs can be ineffective or lead to new phenotypes (*Tosi et al., 2023; Ngawa et al., 2023; Baranello et al., 2024*). This highlights a pressing requirement to deepening our understanding of disease pathogenesis, identify robust molecular biomarkers for disease severity and treatment response, enabling more informed clinical decision-making before and during the therapeutic intervention period.

*SMN1* gene deletions and *SMN2* copy number are routinely used in clinical practice as diagnostic and prognostic biomarkers. Although *SMN2* mRNA and SMN protein levels may help in patient stratification (*Wadman et al., 2016*) they display high heterogeneity (*Signoria et al., 2024*). Similar results were obtained when evaluating therapeutic responses to DMTs (*Poirier et al., 2018; Wirth et al., 2021; Baranello et al., 2021; Wadman et al., 2016; Kolb et al., 2017; Otsuki et al., 2018; Chiriboga et al., 2016; Chiriboga et al., 2023; Mercuri et al., 2025, Baranello al., 2021; Finkel et al., 2017*). Moreover, DMTs do not always lead to an improvement in motor function, likely due to the varying SMN requirements across different tissues and developmental stages (*Groen et al., 2018; Uhlén M et al., 2015*). These findings suggest that *SMN2* mRNAs and SMN protein levels alone may not be reliable indicators for monitoring disease progression or for accurately reflecting therapeutic efficacy in SMA patients (*Wadman et al., 2016; Signoria et al., 2024*). Likewise, various molecules obtained from diverse patients’ biofluids (*Glascock et al., 2023; Maretina et al., 2024; Meneri et al., 2023*) have been proposed as molecular biomarkers for prognosis (*Darras et al., 2019*) and assessment of drug efficacy (*Faravelli et al., 2020; Errico et al., 2022; Maretina et al., 2024*). Among them, neurofilaments (NFLs) are eliciting some interest (*Bayoumy et al., 2024; Faravelli et al., 2020*). However, NFLs do not show consistency across different therapeutic treatments (*Flotats-Bastardas et al., 2023*), and are not disease-specific, being mostly limited to detect general neurodegeneration. Since SMA is increasingly recognized as a more complex, multisystemic disorder (*Yeo and Darras, 2020; Detering et al., 2022*), there is an obvious and urgent need for additional biomarkers to improve patient monitoring and to optimize treatment strategies, especially in the post-therapeutic era with non-neuromuscular phenotypes emerging (*Tosi et al., 2023; Ngawa et al., 2023; Baranello G, 2024*). SMN, while primarily known for its role in ribonucleoparticle (RNP) biogenesis (*Price et al., 2018*), has been shown to associate with ribosomes and polysomes in various cellular and animal models (*Bernabò et al., 2017; Sanchez et al., 2013; Sharma et al., 2024; Lauria et al., 2020; Bechade et al., 1999*). It functions as a ribosome-associated protein (RAP) involved in regulating translation (*Bernabò et al., 2017; Lauria et al., 2020; Susanto et al., 2024*), recruiting a subset of mRNAs (*Lauria et al., 2020*). Through this function, SMN connects translation to multiple cellular pathways disrupted in SMA (*Zhang et al., 2013; Lotti et al., 2012; Simon et al., 2021; Zillo et al., 2022; James et al., 2021; Wishart et al., 2014; Fallini et al., 2016; Deng et al., 2022; Rossol et al., 2003; Hensel et al., 2018; Tapken et al., 2024; Sharma et al., 2024*). Importantly, reduced SMN levels impair translation *in vitro* (*Bernabò et al., 2017; Sanchez et al., 2013; Fallini et al., 2016; Lauria et al., 2020; Deng et al., 2022*) and *in vivo* in SMA mouse models (*Bernabò et al., 2017; Lauria et al., 2020; Tapken et al., 2024*), as evidenced by decreased protein synthesis in neuronal and peripheral tissues (*Bernabò et al., 2017*) and by prenatal proteome alterations in severe SMA mice (*Soltic et al., 2019; Motyl et al., 2020; Lauria et al., 2020*). These findings highlight the potential of dysregulated aspects of translation to serve as a source of translation-based and disease-specific biomarkers to inform and guide SMA therapeutic strategies (*Sharma et al., 2024*).

To explore this hypothesis, we investigated the impact of SMN deficiency on protein translation in brain, spinal cord and liver at different developmental stages of the Smn^2B/-^ mouse model of SMA (*DiDonato et al., 2001; Hammond et al., 2010; Bowerman et al., 2012*). We identified a set of mRNAs undergoing early alterations in ribosome recruitment and validated these results in patient-derived fibroblasts before and after treatment with the *SMN2* splicing modifier Risdiplam. We determined the presence of transcriptional independent and translation-specific defects, which are shared across patients and mouse models, and identified Collagen, type I, alpha 1 (COL1A1) as a novel potential translation-based biomarker for SMA.

## Results

### The Smn^2B/-^ mouse model shows translational alterations across multiple tissues

To identify candidate mRNAs with altered ribosome recruitment suitable for subsequent validation in patient-derived fibroblasts, we first used the Smn^2B/-^ mouse model of SMA (*Hammond et al., 2010; Bowerman et al., 2012*). We reasoned that early translational changes in this model may uncover conserved biomarkers of disease progression. To determine whether translational alterations in this model were consistent with results obtained from the more severe Taiwanese mouse model of SMA (*Bernabò et al., 2017; Lauria et al. 2020*), we mapped the progression of translation defects in brain, spinal cord and liver from different stages of disease progression using polysome and ribosome profiling. We defined pre-symptomatic (P5), early symptomatic (P10), and late symptomatic (P18) stages of the disease, based on changes in body weight (**Supplementary Figure 1A**, *Reilly et al.,2024, Bowerman et al., 2012; Eshraghi et al., 2016).* Consistently, downregulation of SMN protein was confirmed at both early and late stages of disease progression (**Supplementary Figure 1B and 1C**). Following the approach used with the more severe Taiwanese mouse model of SMA base on polysome profiling (*Bernabò et al., 2017*), we quantified the Fraction of Ribosomes in Polysomes (FRP) at the three stages of disease progression (**Figure 1A**).

**Figure 1.**
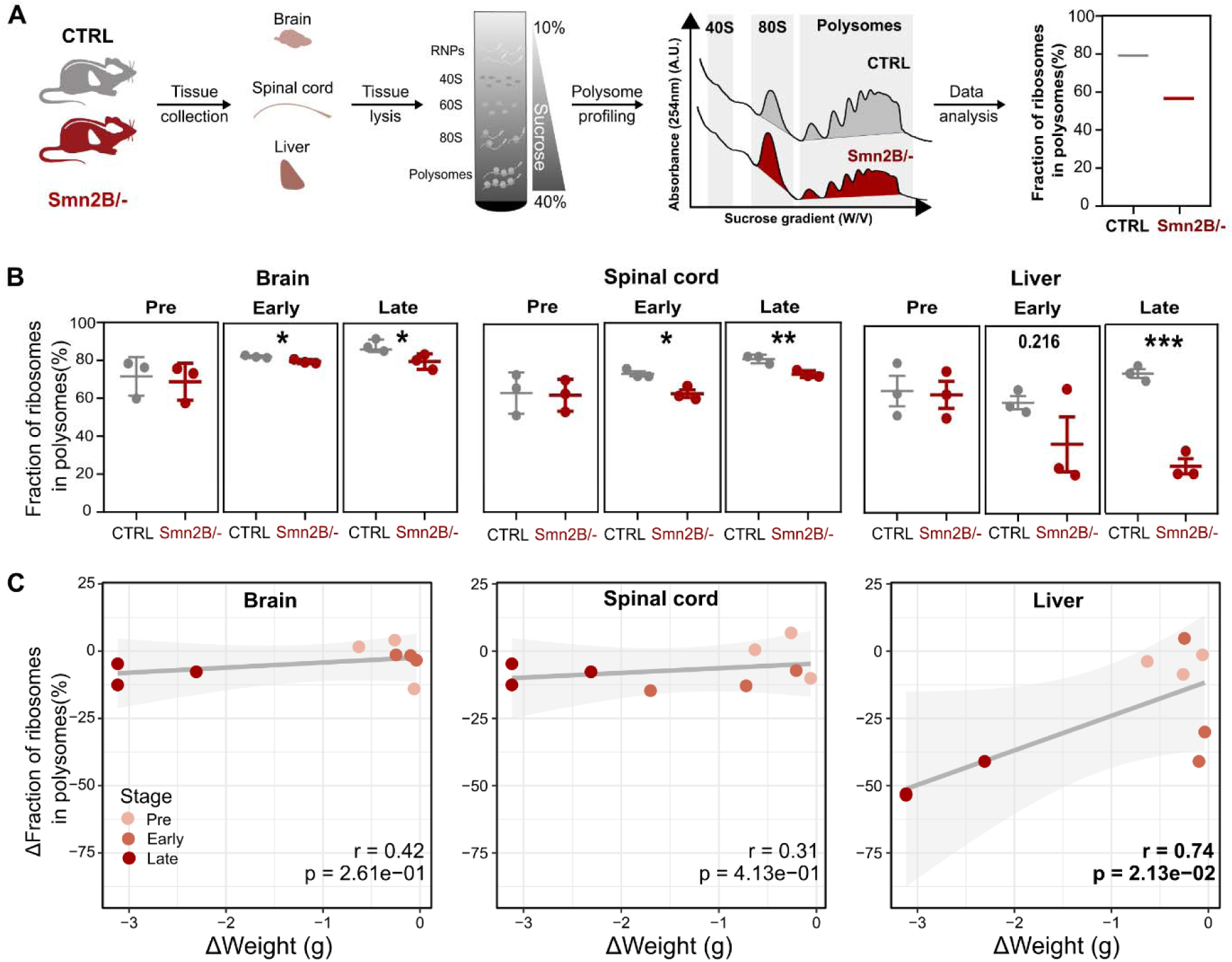
Translation impairment in brain, spinal cord, and liver of Smn^2B/-^ mice compared to littermate controls. **(A)** Schematic representation of the polysome profile from control and SMA tissues to calculate the Fraction of Ribosomes in Polysomes. **(B)** Comparison of Fraction of Ribosomes in Polysomes (FRP) betwee Control and SMA tissues at different stages of disease. Data are meanD±Ds.e.m. of nD=D3 biologically independent experiments. Significance was calculated by unpaired two-tailed Student’s t-test and defined as *p<0.05 and **pD<D0.01. Brain (p-values Early p = 0.04383; Late p = 0.03707), Spinal cord (p-values Early p = 0.01235; Late p = 0 .00845), and Liver (p-value Late p = 0.00044). **(C)** Relationship between the change in bod weight and the corresponding change in FRP. The ΔFraction of ribosomes in polysomes and Δweight are calculate by subtracting the average FRP or weight of control mice from the individual SMA mouse. Pre = pre-symptomatic (P5), Early = Early symptomatic (P10) and Late = Late symptomatic (P18). Regression line is 99% confidence level interval, Pearson correlation coefficient, and its p-value are displayed. Significant p-values<0.05 are highlighted in bold.

The FRP provides an estimate of ribosome engagement in polysomes and of translation efficiency (*Bernabò et al., 2017*). Whilst no significant changes were identified at the pre-symptomatic stage, significant decreases in FRP appeared in brain and spinal cord at both early (P10) and late (P18) symptomatic stages (**Figure 1B**), as previously observed in the Taiwanese mouse model (*Bernabò et al., 2017*). In line with recent reports on liver alterations in this model (*Reilly et al., 2024*), but in striking difference to what was reported in the more severe mouse model of SMA (*Bernabò et al., 2017*), the most substantial decrease in FRP appeared in liver at early and late stages, suggesting more severe translation impairment in this tissue compared to the brain and spinal cord (**Figure 1B**). Importantly, these changes in the FRP are not driven by alterations in the ERK and mTOR pathways that are known to regulate translation (*Sonenberg & Hinnebusch, 2009*) (**Supplementary Figure 1D-F**), and mildly correlate with the decrease in mouse body weight (**Figure 1C**). These findings confirmed that translation defects in the Smn^2B/-^ mouse model of SMA are associated with phenotypic readouts of disease progression and that dysregulation of FRP is a common feature of both the Taiwanese mouse model (*Bernabò et al., 2017*) and the Smn^2B/-^ mouse model of SMA.

### Prioritization of candidate biomarkers through the identification of common translationally-dysregulated processes across multiple tissues in SMA

Prompted by the disease-dependent changes observed in the FRP, we investigated the transcript-specific alterations in ribosome occupancy at genome-wide level by reanalysing ribosome profiling data obtained from early symptomatic spinal cord (*Tapken et al., 2024*), and by performing the same analysis in brain and liver at the early symptomatic stage (P10) (**Figure 2A, Supplementary Figure 2A**). By differential expression analysis, we identified 80, 372, and 1284 differentially translated transcripts (DEGs) of protein coding genes in the brain, spinal cord, and liver, respectively (**Figure 2B, Supplementary Figure 2B, Supplementary File 1**). In accordance with previous results (**Figure 1B**), the largest number of altered transcripts were observed in the liver.

**Figure 2.**
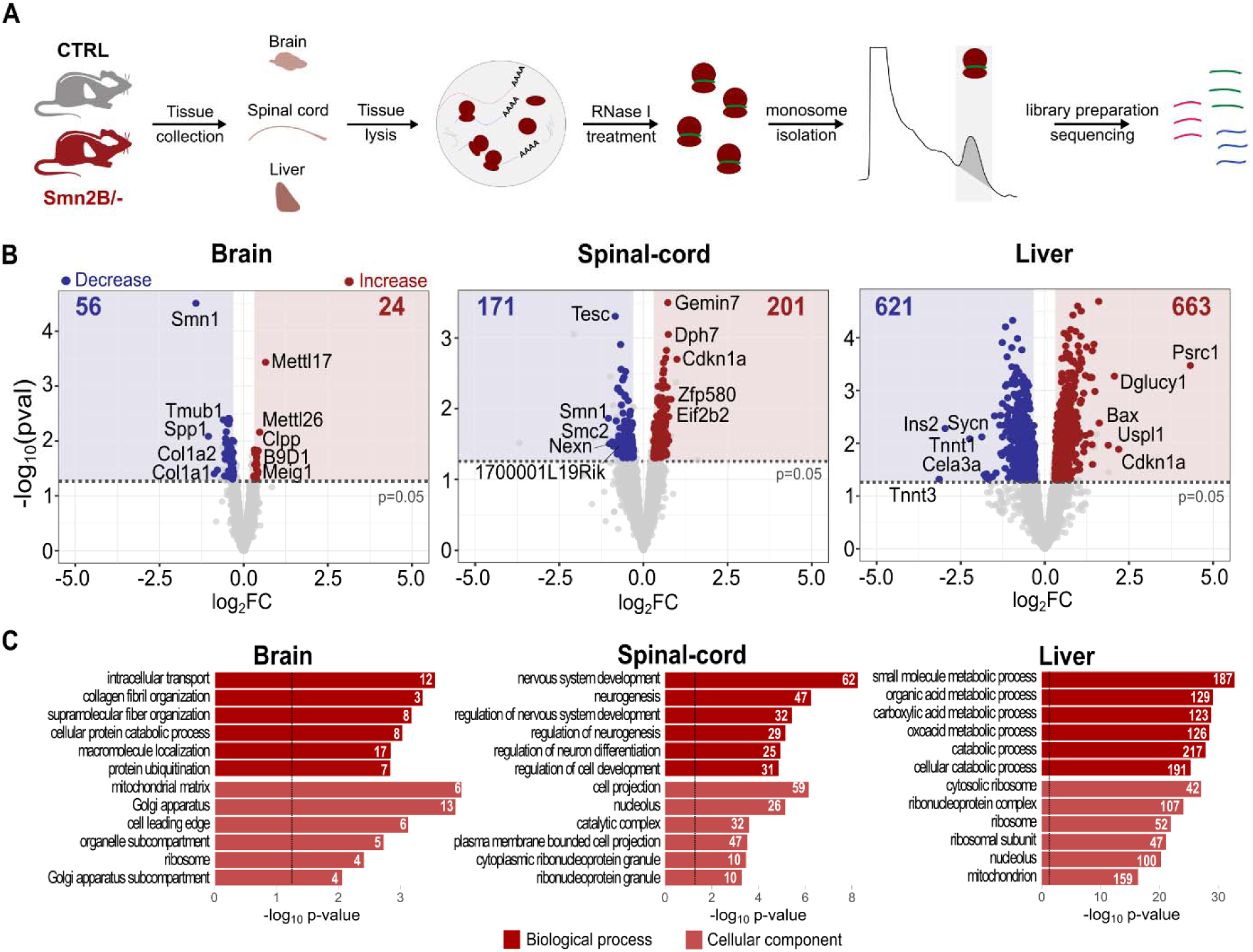
mRNAs exhibit differential ribosome occupancy in Smn^2B/-^ mice compared to littermate controls. **(A)** Schematic representation of ribosome profiling from early symptomatic (P10) Smn^2B/-^ (n=3) and control mice (n=3) tissues. **(B)** Volcano plot showing differentially expressed transcripts. The number of transcripts showing decrease or increase in ribosome occupancy is mentioned in each panel. Names of the top five significantly up and downregulated transcripts in ribosome occupancy are labelled on the plot. Transcripts showing significantly differential ribosome occupancy were defined by |log_2_FC|>0.3, and p-value<0.05. Significance was calculated usin Quasi-likelihood F-test. **(C)** Enrichment analysis of Gene Ontology terms in the brain, spinal cord, and liver. The bar lengths report the significance of the enrichment, and the number of genes associated within each category is reported in each bar. The vertical dotted line represents the p-values.

Not surprisingly, Gene Ontology analysis of these DEGs revealed tissue-specific enrichment of genes: collagen fibril organization in the brain; neurodevelopmental processes in the spinal cord, and metabolic pathways in the liver. Interestingly, the term "Ribosome" was commonly enriched in both brain and liver datasets, highlighting shared ribosome-related alterations, which are consistent with findings in the brain of the Taiwanese SMA mouse model (*Bernabò et al., 2017; Lauria et al., 2020*) (**Figure 2C**). These results demonstrate the presence of widespread alterations in ribosome occupancy in mRNAs with tissue- and model-independent features across brain, spinal cord, and liver in the Smn^2B/-^ mouse model of SMA.

To select candidate biomarkers, we searched for translationally dysregulated transcripts that are shared across multiple tissues in SMA (**Figure 3A**). As observed in previous analysis (*Tapken et al., 2024*), no individual DEG was changing across all tissues (**Figure 3A** and **Supplementary Table 1-3**). We therefore followed a systems approach as in *Tapken et al., 2024* and selected all transcripts that showed a significant change in at least two tissues (a total of 64 transcripts) (**Figure 3B**). Among these, *Cdkn1 and Snrpa1*, transcripts known to be altered in SMA (*Bäumer et al., 2009; Cherry et al., 2017; Corti et al., 2008; Nichterwitz et al., 2020; Olaso et al., 2006; Ruggiu et al., 2012; Tadesse et al., 2007; X. Zhang et al., 2013; Reedich et al., 2021; Doktor et al., 2017*), showed a significant increase in ribosome occupancy in the spinal cord and liver and a trend of increased ribosome occupancy also in the brain. By gene set enrichment analysis of these 64 transcripts (KEGG pathway) (**Figure 3C**), we found that the top three enriched terms were associated with ribosomes, extracellular matrix-receptor interaction, and RNA transport. Similarly, by protein-protein network analysis using the STRING database, we identified four main protein networks associated with ribosome/translation, RNA splicing/transport, extracellular matrix, and DNA replication (**Figure 3D**). The transcripts forming each cluster comprise mRNAs from all three tissues, suggesting the presence of commonly dysregulated processes.

**Figure 3.**
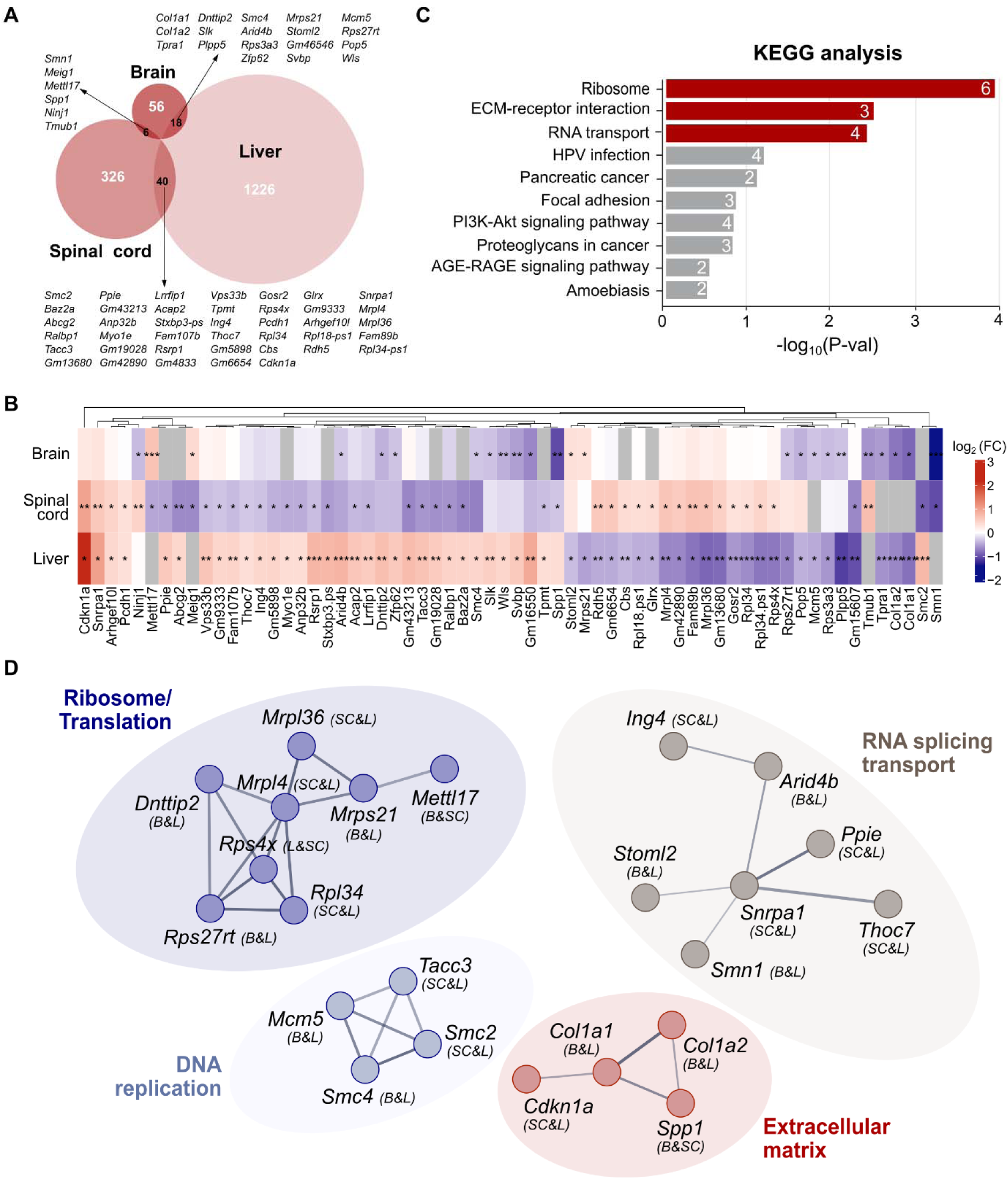
Summary of bioinformatics analysis of shared differentially translated mRNAs in Smn^2B/-^ mouse tissues. **(A)** Venn diagrams illustrating the overlap between significantly dysregulated transcripts in three tissues. Names of the shared genes between all combinations of two tissues are indicated. **(B)** Heatmap detailing the extent of the log_2_ fold changes in shared genes in all three tissues with blue for down and red for upregulated genes. The grey color represents transcript not detected in those tissues. Significant p-values are reported on the plots. * p-val<0.05,** p-val<0.01, and *** p-val<0.001. Significance was calculated using Quasi-likelihood F-test. **(C)** Bar charts showing the top significantly enriched terms in KEGG analysis of shared differentially expressed transcripts. The bar lengths report the significance of the enrichments, and the number of genes associated with each category is reported inside each bar. The red colour represents the significant terms (-log10(p-val) = 1.3). **(D)** Protein-protein network is formed by dysregulated transcripts that overlap between the tissues, as shown in (A). The 4 clusters, which are formed by the contribution of DEGs for all three tissues, are: (i) Ribosome/translation formed by Dhttip2 (B & L), Mrpl36 (SC & L), Mrpl14 (SC & L), Mrps21 (B & L), Rps4x (SC & L), Rpl34 (SC & L), Rps27rt (B & L) and Mettl17 (B & SC); (ii) RNA Splicing/transport: Ing4 (SC & L), Arid4b (B & L), Snrpa1 (SC & L), Stoml2 (B & L), Smn1 (B & SC), Ppie (SC & L) and Thoc7 (SC & L); (iii) DNA Replication: Mcm5 (B & L), Smc4 (B & L), Smc2 (SC & L), Tacc3 (SC & L); (iv) Extracellular matrix: Col1a1 (B & L), Col1a2 (B & L), Cdkn1a (SC & L) and Spp1 (B & SC). B stands for brain, SC for spinal cord and L for liver.

Interestingly, COL1A1 and COL1A2 proteins have been identified as differentially expressed proteins in previous proteomic studies on fibroblasts and in the cerebrospinal fluid (CSF) from SMA patients with milder forms of the disease (*Brown et al., 2022; Kessler et al., 2020*). Moreover, *Col1a1* is part of the collagen-related mRNAs that are regulated by SMN-primed ribosomes (*Lauria et al., 2020*). Thus, three mRNAs from the extracellular matrix group (*Col1a1*, *Col1a2*, and *Spp1*) were shortlisted as potential translation-specific biomarkers.

### Early symptomatic translation- and disease-specific changes in *Col1a1*

To validate that the observed alterations of *Col1a1*, *Col1a2*, and *Spp1* were translation -specific and transcription-independent, according to the role played by SMN in translation (*Lauria et al., 2020; Sharma et al., 2024*), we investigated the changes in cytoplasmic mRNA levels at pre-symptomatic (P5), early symptomatic (P10), and late symptomatic (P18) stages in the brain, spinal cord, and liver of Smn^2B/-^ mice compared to littermate controls. In all cases we did not observe any significant change in transcript levels, although there were changes in *Spp1* in the liver at late symptomatic stages of disease (P18) (**Figure 4A** and **Supplementary Figure 3A**). These results support the hypothesis that alterations observed in ribosome profiling of P10 Smn^2B/-^ mice are translation-specific and not caused by changes in mRNA levels.

**Figure 4.**
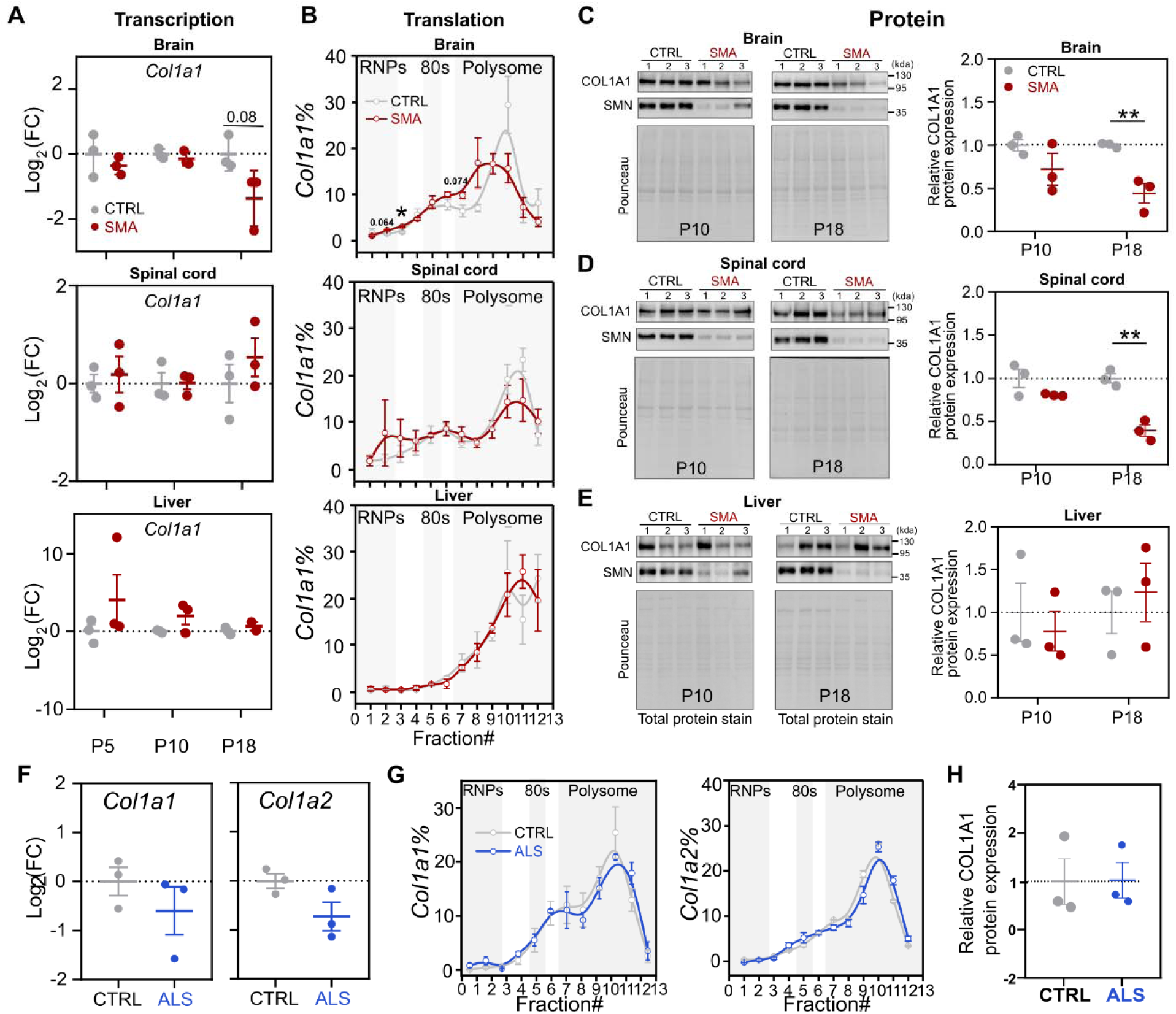
Translational alterations of Col1a1 are transcriptional-independent and disease-specific. **(A)** Quantification of qPCR analysis performed for the *Col1a1* transcripts in cytoplasmic mRNAs from control and SMA brain, spinal cord and liver at the P5, P10, and P18 stages of SMA disease in Smn^2B/-^ mice. Data are mean ± s.e.m. among n = 3 biologically independent samples. Significant changes were assessed using unpaired two-sided t-tests. **(B)** Relative co-sedimentation profile of *Col1a1* mRNAs in control (grey) and SMA (red) brain, spinal cor and liver at P10. Data are mean ± s.e.m. among n = 3 biologically independent samples. Significant changes were assessed using unpaired two-sided Student’s t-tests. * p-val<0.05. Brain (p-value fraction 3 p = 0.01101). **(C-E)** Protein levels of Col1a1 and SMN in the brain (C), spinal cord (D), and liver (E) at P10 and P18 in controls and SMA using western blot analysis. Quantification of immunoblots for Col1a1 is normalized to total protein stain. SMA expression values were normalized and compared with control values for each of the tissues. Data are mean ± s.e.m. among n = 3 biologically independent samples. Significant changes were assessed using unpaired two-side Student’s t-tests. * p-val<0.05 and ** p-val<0.01. Brain (p-value P18 p = 0.00794), Spinal cord (p-value P18 p =0.00219). **(F,G)** Quantification of qPCR analysis for *Col1a1* and *Col1a2* (F) relative co-sedimentation profile of *Col1a1* and *Col1a2* mRNAs and protein levels of Col1a1 (G) in the spinal cord and cortex tissues of a mouse model of Amyotrophic Lateral Sclerosis (blue) and littermate controls (grey). (**H**). Data are mean ± s.e.m. among n = 3 biologically independent samples. Significant changes were assessed using unpaired two-sided Student’s t-tests.

Next, using a co-sedimentation analysis of mRNAs with polysomes to evaluate the engagement of ribosomes with mRNAs (*Lauria et al., 2020*), we measured the relative distribution of the three mRNAs along the sucrose gradient fractions of polysome profiling from control and early symptomatic mouse brain, spinal cord, and liver. *Tuba4a* mRNA was chosen as a control as it is not a SMN-specific mRNA (*Lauria et al., 2020*), and showed no difference in co-sedimentation in either brain or spinal cord (**Supplementary Figure 3B-C**, top panel) and a very mild, but significant, decrease in the ribosome (80S) fraction at the early symptomatic stage of SMA in the liver (**Supplementary Figure 3D** top panel).

In brain at the early symptomatic (P10) stage of disease, *Col1a1* (**Figure 4B**, top panel) *and Col1a2 (***Supplementary Figure 3B**, middle panels) shifted from polysomal fractions towards free RNA (RiboNucleoParticles (RNPs)) fractions. A similar trend was observed for *Col1a1* in spinal cord (**Figure 4B**, middle panel), whilst no major changes in *Col1a1* (**Figure 4B**, lower panel) and *Col1a2* (**Supplementary Figure 3D**, middle panel) were observed in the liver. *Spp1* showed a mild, non-significant shift towards lighter fractions only in the liver (**Supplementary Figure 3B-D**, lower panels). Importantly, in brain and spinal cord the reorganization of ribosomes on *Col1a1* in brain and spinal cord reflect a later decrease in COL1A1 protein levels at the P18 stage compared to age-matched controls (**Figure 4C** and **4D**, right panels), while no change was identified in the liver (**Figure 4E**, right panel). These results demonstrate that *Col1a1* exhibits translation-specific and transcription-independent alterations, which lead to changes in protein levels which are significant at late symptomatic stages of disease.

To understand if the identified translation alterations of *Col1a1* are disease-specific, we performed a similar analysis in a mouse model of Amyotrophic Lateral Sclerosis (ALS, TDP-43^Q311K^) (*Arnold et al., 2013*), a neuromuscular disorder closely related to SMA. In cortex and spinal cord from pre-symptomatic (3-month-old) animals, no significant changes were observed either in mRNA levels or in the co-sedimentation profile for *Col1a1* or *Col1a2* (**Figure 4F** and **G**). Protein levels of COL1A1 did also not change (**Figure 4H** and **Supplementary Figure 3E**), suggesting that the observed alterations in *Col1a1* translation in SMA are also disease specific.

### Clinical relevance of translational defects in *Col1a1* in SMA

To further explore the clinical potential of *Col1a1* as a translation- and disease-specific biomarker, we used patient-derived fibroblasts obtained from paediatric and adult SMA patients (**Supplementary Table 4**). These fibroblasts have been obtained from two different sources (the Coriell Institute, US and University Medical Center Utrecht UMCU, NL) and have recently been shown to be reliable models of SMA as they recapitulate both heterogeneity and complexity of patients’ response to therapies (Signoria et al., 2024).

First, we analysed mRNA levels, translation, and protein levels of COL1A1 in the Coriell cohort of fibroblasts (three healthy donors, one carrier, and four SMA patients (Type I or Type II)) (**Figure 5A**). As observed in mouse tissues, no significant changes were observed in the overall *COL1A1* transcript levels (**Figure 5A**, left panel). Next, using polysome profiling (**Supplementary Figure 4A**), we performed a co-sedimentation analysis of *COL1A1* mRNAs and compared its distribution in healthy and SMA fibroblast lines (**Figure 5A**, middle panels, and **Supplementary Figure 4B)**. In paediatric SMA Type II and Type I fibroblasts, the *COL1A1* mRNA shifted from polysomes towards lighter fractions, with similar trends to those observed in mouse tissues.

**Figure 5.**
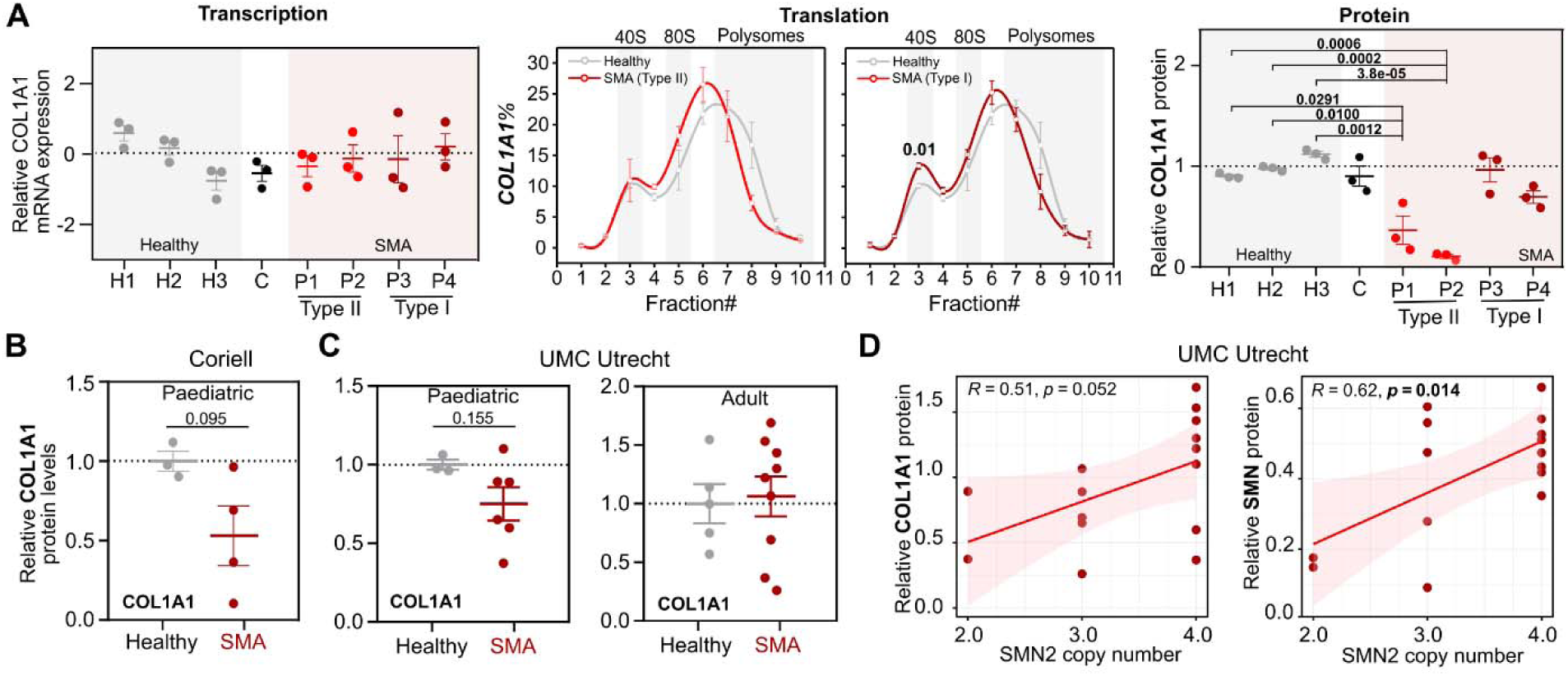
COL1A1 translational specific downregulation in patient derived fibroblasts. **(A)** *COL1A1* mRNA levels in total RNA from healthy, carrier (H1-3 and C) and SMA patient-derived fibroblasts (P1-4, left). Data are mean ± s.e.m. among n = 3 independent technical replicates for each individual. Comparison between relative co-sedimentation profiles of *COL1A1* mRNA from healthy (n=3) and SMA patient-derived fibroblasts SMA Type I (n = 2), SMA Type II (n = 2) (middle). Data are mean ± s.e.m. among n biological independent replicates. COL1A1 protein expression in healthy, carrier and SMA patient-derived fibroblasts (left), data are mean ± s.e.m. among n = 3 independent technical replicates. Expression values were normalized and compared with healthy control values. Significant changes between each healthy sample and the other samples were assessed using one-way ANOVA test, with Holm correction of p-values. Significant p-values are reported on the plots. **(B, C)** Comparison of COL1A1 protein expression in Coriell Institute cohort (Healthy n=3, SMA n=4) and UMCU cohort-paediatric (Healthy n=3, SMA n=6) and adult (Healthy n=5, SMA n=9). Data represent the mean ± s.e.m. of n individual, shown as a dot, for each group. Significant changes were assessed using unpaired two-sided Student’s t-tests. **(D)** Corelation between COL1A1/SMN protein expression and SMN2 copy number in SMA patients’ fibroblasts from the UMCU cohort. Data are plotted as mean of n=3 independent technical replicates. The regression line, its 99% confidence level interval, the Pearson correlation coefficient, and its p-value are displayed. Significant p-value<0.05 are highlighted in bold.

Clustering analysis of the percentage of *COL1A1* mRNA in each fraction showed that co-sedimentation profiles of SMA Type II and Type I fibroblasts grouped together, distinguishing them from healthy controls (**Supplementary Figure 4C**). These translational alterations correspond to a substantial reduction in COL1A1 protein levels in the two SMA Type II fibroblasts compared to healthy and carrier samples (**Figures 5A**, right panel, **Figure 5B** and **Supplementary 4D**). Our findings indicate a significant reduction in COL1A1 protein levels in SMA Type II patient fibroblasts from the Coriell Institute cohort. This decrease likely results from translational dysregulation and is independent of transcript abundance.

To strengthen our findings, we analysed COL1A1 protein levels in the second cohort from UMCU. In this case we investigated both paediatric and adult SMA fibroblasts (**Figure 5C**). A trend of decrease in COL1A1 levels was observed in the paediatric SMA samples (**Figure 5C**, right panel), consistent with results from the Coriell cohort. However, no changes were detected in the adult samples **(Figure 5C**, middle panel**, Supplementary Figures 5 and 6)**. Interestingly, in SMA patients COL1A1 protein expression correlates positively with the *SMN2* copy number and SMN protein expression (**Figure 5D**), suggesting that COL1A1 expression may be dependent on SMN protein levels, opening the possibility that it may serve as a biomarker for SMN-restoring interventions.

To explore this possibility, we treated the UMCU fibroblasts with the *SMN2* splicing modifier Risdiplam (**Figure 6A**) and assessed SMN (**Figure 6B, Supplementary Figure 5**) and COL1A1 protein levels (**Figure 6C, Supplementary Figure 6**). By grouping fibroblasts in healthy controls, paediatric and adult SMA, we observed that Risdiplam treatment led to a significant increase in SMN level only in fibroblasts derived from adult SMA patients (**Supplementary Figure 5**). In contrast, COL1A1 expression levels remained unchanged across all groups (**Supplementary Figure 6**). When analysing individual responses to treatment, all paediatric (3/3) and 6 out of 7 adult SMA-derived fibroblasts showed a significant increase in SMN level upon treatment. The remaining adult SMA (1/7) and all healthy donor fibroblasts (3/3) showed no significant change in SMN expression (**Figure 6D**). Regarding COL1A1, strikingly, all paediatric SMA fibroblasts (3/3) and one adult (1/7) SMA fibroblast exhibited a significant increase in protein level. On the contrary, the remaining adult SMA lines (6/7) showed a downregulation, with 4 reaching statistical significance (**Figure 6E**). Similarly, healthy donor fibroblasts (3/3) showed a trend of downregulation, with two lines showing a statistically significant decrease (**Figure 6E**).

**Figure 6.**
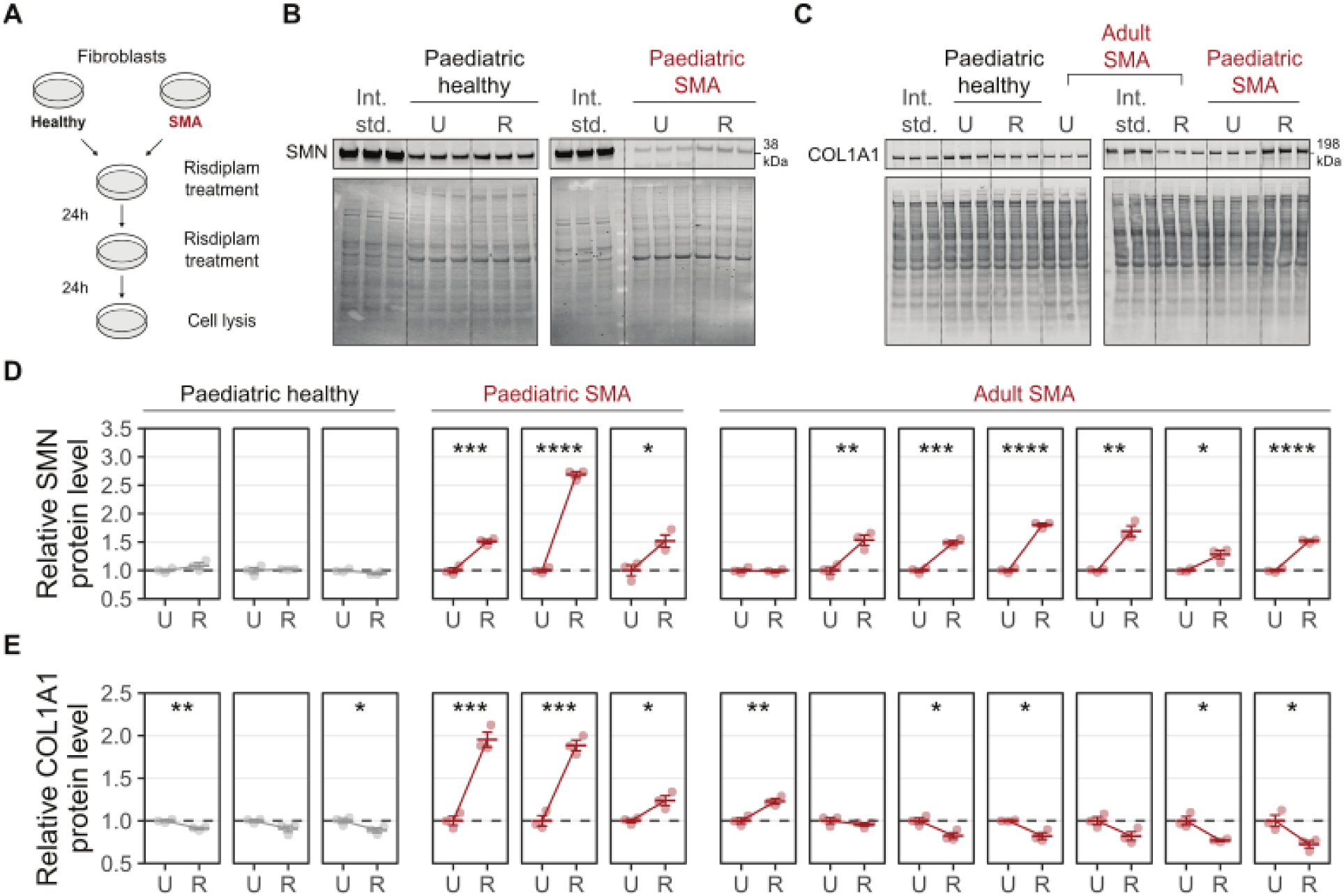
Risdiplam rescues COL1A1 protein expression in fibroblasts derived from paediatric SMA patients. **(A)** Schematic representation of the Risdiplam treatment. **(B)** Representative immunoblots for SMN in untreated (U) and Risdiplam-treated (R) patient-derived fibroblasts from 3 technical replicates of one paediatric patient and one healthy donor. The internal standard (Int. std.) is used to compare protein signal across different membranes. Each blot is accompanied by a corresponding total protein stain (lower panel). **(C)** Representative immunoblots for COL1A1 in untreated (U) and Risdiplam-treated (R) patient-derived fibroblasts from paediatric healthy donors, and paediatric and adult SMA patients. The internal standard (Int. std.) is used to compare protein signal across different membranes. Each blot is accompanied by a corresponding total protein stain (lower panel). **(D)** Quantification of SMN protein level of Risdiplam-treated (R) relative to the untreated (U) fibroblasts. Protein levels were normalize for total protein stain and internal standard. Data are mean ± s.e.m. among n = 3 technical independent replicates. Significant changes were assessed using unpaired two-sided Student’s t-test, with Holm correction of p-values. * p-val<0.05, ** p-val<0.01, *** p-val<0.001, and **** p-val<0.0001. **(E)** Quantification of COL1A1 protein level of Risdiplam-treated (R) relative to the untreated (U) fibroblasts. Protein levels were normalized for total protein stai and internal standard. Data are mean ± s.e.m. among n = 3 technical independent replicates. Significant changes were assessed using unpaired two-sided Student’s t-test, with Holm correction of p-values. * p-val<0.05, ** p-val<0.01, *** p-val<0.001, and **** p-val<0.0001.

Overall, these findings suggest that COL1A1 is a promising biomarker for SMA that correlates with disease severity as well as response to DMT in paediatric SMA patients.

## Discussion

The significant advances achieved by SMN-restoration therapies (*Finkel et al., 2017; Mendell et al., 2017; Poirer et al., 2018*) are producing a substantial change in SMA management (*Moultrie et al., 2025; Yeo et al., 2024*). With the three SMN-dependent therapies approved for treating individuals living or being prenatally diagnosed with SMA, it is likely that soon almost every patient will be treated at some stage of disease (*Govoni et al., 2018; Motyl and Gillingwater 2022*). While lifespan and motor abilities may improve following treatment (albeit by variable amounts), additional, and often unforeseeable, complications and comorbidities are reported (*Westrate et al., 2022; Baranello et al., 2024*). Thus, current SMN restoration therapies remain only partially effective and do not represent a definitive ‘cure’ for SMA (*Chaytow et al., 2021*).

Considering the notable but heterogeneous impact of these innovative therapies, it is essential to develop methods for detecting treatment-induced changes in disease characteristics and thereby discover a new generation of biomarkers (*Yeo et al., 2024*). Currently proposed biomarkers fall into two large categories: biomolecular and physiological/clinically assessed indicators (*Pino et al., 2021; Meneri et al., 2023*). With few exceptions, most of these biomarkers are SMA-unspecific, i.e. not based on molecular mechanisms directly related to SMA. In addition, they largely represent a neuromuscular-related readout of disease progression or therapeutic efficacy. Even neuromuscular-related biomarkers that are eliciting excitement, such as neurofilaments (p-NfH, or NfH), seem to show quite limited dynamic range of activity and are largely disease-unspecific (*Wurster et al., 2019; Faravelli et al., 2020*). Therefore, at present, no reliable and SMA-specific molecular biomarker, not even the number of *SMN2* copies (*Wadman et al., 2020*), can be employed to robustly monitor disease severity and progression, or response to specific treatments, suggesting that we still need to grasp the full spectrum of SMN functions to facilitate the identification of valid biomarkers.

In this work, we explored the hypothesis that SMN-dependent alterations in translation observed in SMA models (*Bernabò et al., 2017; Lauria et al., 2020; Tapken et al., 2024; Sharma et al., 2024*) may represent a still understudied source of a new generation of biomarkers. Utilising multiple techniques, we observed a significant reduction in translation at early and late symptomatic stages across the brain, spinal cord, and liver of the Smn^2B/-^ mouse model (*Bowerman et al., 2012*). This indicates that impaired translation persists throughout disease progression as observed at early and late stages of SMA across brain and spinal cord in the severe Taiwanese mouse model (*Bernabò et al., 2017; Lauria et al., 2020*). These findings suggest a common downregulation of translation in the brain and spinal cord across different SMA mouse models. Interestingly, the liver shows robust translational alterations in the Smn^2B/-^ model. Consistent with our findings, liver-specific defects have also been observed in Smn^2B/-^ mice prior to the onset of neuronal pathology, suggesting a potential early metabolic vulnerability in SMA (*Reilly et al., 2024; Almeida et al 2025*). The strong correlation between the decrease in translation in liver and body weight loss in Smn^2B/-^ mice underscores the relevance of translational defects and opens a new hypothesis as to what mechanism underlies the liver dysfunction observed in this tissue (*Deguise et al., 2021; Szuyogova et al., 2016*) and in patient-derived hepatocytes (*Leow et al., 2024*).

Through a translatome analysis of the of early symptomatic tissues, we showed the genome-wide effect of translational defects caused by SMN deficiency in the Smn^2B/-^ model and revealed hundreds of mRNAs exhibiting significantly altered ribosome occupancy across tissues. A significant overlap of DEGs from brain and spinal cord, spinal cord and liver, and liver and brain led to shortlisting three potential translation-based and disease-specific biomarkers, which are part of the extracellular matrix (ECM) network (*Col1a1*, *Col1a2*, and *Spp1*). Among these three mRNAs, *Col1a1* was identified as an “SMN-specific” mRNA (*Lauria et al., 2020*).

Strikingly, despite no significant change at the transcriptional level, a strong depletion of *Col1a1* mRNA from polysomes in brain and spinal cord confirms its defective translation and protein levels in SMA. The absence of changes in the liver are likely due to *Col1a1* low expression level in this tissue (*Uhlén et al., 2015*), compared to brain and spinal cord. Furthermore, we confirmed that the translation defect of *Col1a1* is SMA-specific. In fact, tissues from pre-symptomatic ALS-affected mice showed no changes in *Col1a1* translation. The role of COL1A1 in disease pathogenesis may be relevant in the framework of SMA as a multisystemic disorder, as shown by the collagen dysregulation in the Taiwanese mouse model of SMA, in which type IV collagen protein alterations were observed in the kidney (*Allardyce et al., 2020*). Likewise, mRNA-level changes in *Col12a1*, *Col5a2*, and associated ECM remodelling factors (*Lox*, *Ogn*, *Fmod*) were detected in bone tissue (*Hensel et al., 2020*), altogether suggesting a widespread developmental ECM deficiency in SMA mouse models. In line with these results, mutations/deletions in the *COL1A1* gene in humans cause alterations in bone and vascular systems and in and many other tissues (*Fajardo-Jiménez et al., 2022*). Interestingly, defective vascular system (*Somers et al., 2016; Zhou et al., 2022*), weakened bones, and frequent fractures are prevalent in SMA patients (*Baranello et al., 2019; Khatri et al., 2008; Vai et al., 2015; Wasserman et al., 2017*).

To determine the clinical relevance of our findings, we utilised fibroblasts from two centres and derived from adult and paediatric patients and analysed COL1A1 levels before and after treatment with the SMN-restoring and SMN splicing modifier Risdiplam (*Poirer et al., 2018*). As in mouse tissues, these fibroblasts exhibited transcription-independent depletion of *COL1A1* mRNA from polysomes leading to a significant decrease in COL1A1 protein expression in paediatric samples. Consistent with our findings, recent proteomics studies have also identified dysregulation of COL1A1 protein in fibroblasts from milder SMA patients (*Brown et al., 2022; Kessler et al., 2020*). Importantly, Risdiplam treatment of paediatric SMA fibroblasts led to an increase in COL1A1 level, underscoring the clinical relevance of our findings and highlighting the potential of *COL1A1* to act as a translation- and SMA-specific biomarker. These findings warrant further investigation into *COL1A1* utility for monitoring disease progression and predicting individual responses to therapy in the emerging ’post-treatment’ era of SMA.

## Materials and Methods

### Smn^2B/-^ mouse model of Spinal Muscular Atrophy

All animal procedures were performed in accordance with the United Kingdom Home Office regulations (Guidance on the Operation of Animals (Scientific Procedures Act) 1986) and institutional requirements. The ‘Smn^2B/-^’ mouse model of SMA was established by breeding the *Smn2B/2B* allele backcrossed onto a C57BL6 background and bred with *Smn+/-* mice obtained from Jackson Laboratories (*DiDonato et al., 2001; Bowerman et al., 2012; Eshraghi et al., 2016*). Phenotypically normal littermates (Smn^2B/+^) were used as controls. Both sexes were used in these experiments. Mice were genotyped using standard protocols. All mice were housed within the animal care facilities in Edinburgh under standard SPF conditions. Based on the onset of SMA symptoms, we classify P5 as pre-symptomatic, P10 as early symptomatic, and P18 as late symptomatic stages, during which tissue samples were collected. All tissues were quickly dissected, snap-frozen, and stored at −80°C until use.

### TDP43 Q331K mouse model of Amyotrophic Lateral Sclerosis

Validation of *Col1a1* and *Col1a2* in another model than the Smn^2B/-^ mouse was performed on cortexes and spinal cords obtained from the pre-symptomatic 3-month-old PrP-hTDP43 Q331K mouse model of Amyotrophic Lateral Sclerosis (ALS) (*Arnold et al., 2013*). In this model, the human TDP-43^Q331K^ has a myc-tag at its N-terminal and is expressed in the central nervous system under the murine PrP promoter (*Arnold et al., 2013*). Mice were purchased from the Jackson Lab (JAX stock #017933) and housed at the Animal Facility at Department CIBIO (University of Trento, Italy). Spinal cord and cortexes from 3-month-old TDP-43^Q331K^ mice and non-transgenic C57BL6/J controls were quickly dissected, snap-frozen, and stored at -80°C until use.

### SMA patient-derived fibroblast cell lines from Coriell Institute of Medical Research

SMA patient-derived fibroblasts were purchased from the Coriell Institute for Medical Science, USA. Cells were grown in a humidified incubator at 37°C with 5% CO_2_ and maintained in Dulbecco’s Modified Eagle Medium high glucose (Euroclone) supplemented with 15% non-inactivated fetal bovine serum (Euroclone), 2 mM L-Glutamine (Euroclone) and Penicillin-Streptomycin solution (Euroclone). For RNA and protein purification, SMA patient fibroblasts were plated at a seeding density of 50,000 cells per well in 12-well plates or 100,000 cells per well in 6-well plates.

### Ethical approval and SMA patients derived fibroblast cell lines from the University Medical Centre Utrecht

Patients included for analysis in this study participate in a prospective, population-based study on SMA in the Netherlands. This study was approved by the UMC Utrecht Medical Ethical Committee (no. 09– 307/NL29692.041.09). We obtained additional written and oral informed consent for skin biopsies from each adult patient and from both parents of each participating minor.

### Study population and genetics

Patient characteristics were collected during interviews with parents using standardized questionnaires and physical examination as part of our ongoing population-based study. Patients were classified as SMA types 1–4 based on the highest achieved motor milestone, following the international SMA classification system with some relevant additions. For all the subjects included, *SMN1* and *SMN2* copy number were confirmed using the SALSA multiplex ligation-dependent probe amplification (MLPA) kit P021 version B1 (MRC Holland) according to the manufacturer’s protocol and Coffyalyser.Net software (MRC Holland).

### Culture of primary fibroblasts from University Medical Centre Utrecht

Primary fibroblasts were obtained from explants of 3-mm dermal biopsies. After 4–6 weeks, fibroblast outgrowths from the explants were enzymatically passaged (Accutase, Sigma-Aldrich, A6964). Fibroblasts were cultured in DMEM containing 4.5 g/L glucose, L-glutamine, and pyruvate (Gibco, 41966-029) with 10% fetal bovine serum (Cytvia, SH30073.03) and 1% penicillin-streptomycin (Sigma-Aldrich, 0000165820). All cell lines were monitored and negative for mycoplasma (Merk, MP0035). For Risdiplam (Sanbio, 29028-1) treatment, cells were treated at a final concentration of 0.5 μM and the treatment was repeated after 24 h. The total treatment time was 48 h.

### Western Blot - primary fibroblasts from University Medical Centre Utrecht

For SMN, semi-quantitative immuno blotting was conducted as previously described (*Signoria et al., 2024; Groen et al., 2018*). For Col1A1, modifications to the protocol were implemented as follows: protein samples were not subjected to heat denaturation. A total of 20 μg of protein was loaded onto a Bolt Bis-Tris Plus Mini Protein 8% gel (Invitrogen, NW00087BOX) and run for 60 minutes at 100V using Bolt MES-SDS running buffer (Invitrogen, B0002). Proteins were subsequently transferred onto a nitrocellulose membrane. Membranes were blocked for 3h in 3% milk. Western blotting was performed using primary antibodies against Col1A1 (Proteintech 67288-1-Ig, 1:2500) and SMN (mouse anti-SMN, BD Biosciences 610647, 1:1,000).

### Sex as a biological variable

Our study examined male and female fibroblasts, and similar findings are reported for both sexes.

### Polysome profiling

For sucrose gradient preparation, solutions were prepared using a unique buffer composition (10 mM Tris-HCl pH 7.5, 10 mM MgCl_2_, 10 mM NaCl) with a variable concentration of sucrose, depending on the gradient type. Normal gradients (13.2 mL) were prepared by overlaying 5.5 mL 40% (w/V) sucrose buffer and filling the tube with the 10% (w/V) sucrose buffer. Small gradients (4 mL) were prepared by overlaying 1.6 mL 40% (w/V) sucrose buffer and filling the tube with the 10% (w/V) sucrose buffer. The gradient was formed by keeping the tube horizontally for 120 min at 4°C.

Cytoplasmic lysates were prepared from mouse-frozen tissues and from patient-derived fibroblasts following the protocol described in *Bernabò et al., 2017*. Briefly, frozen tissues were pulverized in liquid nitrogen using a pestle and a mortar. After pulverization, 800-400 μL of tissue polysome lysis buffer (*Bernabò et al., 2017*) was used for powder resuspension. Fibroblasts were lysed in lysis buffer: 10 mM Tris–HCl pH 7.5, 10 mM MgCl_2_, 10 mM NaCl, 1% w/V Triton X-100, 0.005 U/μL DNAse I, 0.2 U/μL RiboLock RNase inhibitor, 1 mM DTT, 10 μg/mL cycloheximide, 1% w/V Na-deoxycholate. Lysates were kept for 17-20 min on ice to ensure cell lysis. Then, mouse-tissue and patient derived fibroblasts lysates were centrifuged twice or once, respectively, at 12,000 rpm in Eppendorf Centrifuge 5417 for 10 min and the supernatant was collected and loaded on 13.2 mL (mouse tissues) or 4ml (patient-derived fibroblasts) polyallomer ultracentrifuge tubes (Beckman) containing 10%-40% (w/V) linear sucrose gradient. The 13.2 mL gradients were ultracentrifuged for 1 h 30 min at 40,000 rpm at 4 °C in Beckman Optima XPN-100 Ultracentrifuge in SW41 rotor, while 4 mL tubes were ultracentrifuged for 1 h 30 min at 43,800 rpm at 4 °C in Beckman Optima XPN-100 Ultracentrifuge in SW60 rotor. To collect sucrose fractions and measure the absorbance at 254 nm, a Teledyne Isco model 160 gradient analyzer equipped with a UA-6 UV/VIS detector was employed. For 13.2 mL tubes, 12 fractions of 1mL were collected, while for 4 mL tubes 10 fractions of 500 µL were collected. Plotting absorbance vs fraction number yields a polysome profile. The fraction of ribosomes in polysomes (FRP) is calculated using equation:

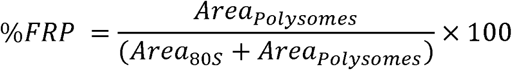

### Co-sedimentation of mRNA with the translation machinery

To analyse the co-sedimentation profiles of mRNAs, a polysomal profiling protocol optimized for tissue and fibroblast analysis was used. The RNA was extracted from each sucrose fraction using the acidic-phenol/chloroform method as described in Lauria et al., 2020. After DNase I treatment, a reverse transcription reaction was performed using the same volume of RNA obtained for each fraction (5 μL) using the RevertAid First Strand cDNA synthesis kit (Thermo Fisher Scientific). qPCR was performed using the CFX Connect Real-Time PCR Detection System (BioRad) using the Kapa Syber Fast qPCR Mastermix (Kapa Biosystems) or qPCRBIO SyGreen Mix Separate-ROX (PCRBiosystem) using primers (**Supplementary Table 5**). The reaction conditions were the following: one step of 95°C for 3 min and 40 cycles of 95°C for 2 s for denaturation, 60-64 °C for 25s for the annealing and elongation phase. The percentage of each transcript distribution along the profile was obtained using the following formula:

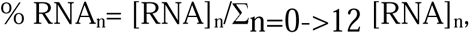

where n is the fraction number, % RNA_n_ is the percentage of the RNA in the fraction n; [RNA]_n_ corresponds to 2(40-Ct) in the fraction n, and 12 is the total number of fractions.

### RT-qPCR on cytoplasmic RNA for differential gene expression

After sucrose gradient fractionation, cytoplasmic RNA was isolated from pooled fractions (1/6^th^ volume from each fraction) using the Acidic-Phenol/Chloroform protocol described in 2020 study (*Lauria et al., 2020*).

The 500 ng total RNA was treated with DNase I and reverse-transcribed using RNA using RevertAid First Strand cDNA synthesis kit (Thermo Fisher Scientific). The RT-qPCR was performed with 1 μL cDNA for each sample. qPCR was performed with qPCRBIO SyGreen Mix separate-ROX (PCRBiosystem) using the CFX Connect Real-Time PCR Detection System (BioRad) using the primers (**Supplementary Table 5**). qPCR conditions were the same as described previously.

The differential expression analysis on total mRNA in mouse tissues has been performed by using *Tuba4a* as a normalizer and the ΔΔCT method, as follows:

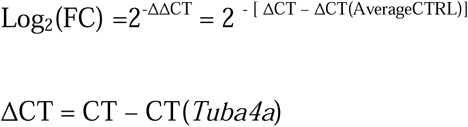

### Total protein extraction and western blotting for mouse tissues and patient’s derived fibroblasts from Coriell Institute of Medical Research

Frozen tissues from both Smn^2B/-^ mice and control littermates and PrP-hTDP43 Q331K mouse model of amyotrophic lateral sclerosis (ALS) were pulverized in liquid nitrogen using a pestle and a mortar. A volume of 30 μL of powder was lysed in 150 μL of RIPA buffer containing 50 mM Tris-HCl pH 8.0, 150 mM NaCl, 1% IGEPAL, 0.5% Na-deoxycholate, and 2% SDS. Lysates were homogenized with the help of the Kimble 749540-0000 Pellet Pestle Cordless Motor and incubated on ice for 15 min. The lysate was then centrifuged at 15,000 rpm in Eppendorf F-45-30-11 Rotor and Eppendorf Centrifuge 5417R at 4°C for 15 min to discard cell debris. The supernatant was transferred into a new tube and stored at -80°C. For fibroblasts, approximately 7*10^5 cells were plated in a six well-plate and left growing until reaching 80 % confluency. Each well is washed 3x with cold PBS. Fifty µL of RIPA lysis buffer (Thermo Scientific) supplemented with: 1 µg/mL pepstatin A, 5 µl/mL Phosphatase Inhibitor Cocktail 2, 5 µL/ml Phosphatase Inhibitor Cocktail 3, 2.5 µL/mL PIC is added to each well and scraped. Lysates were collected and freeze-thaw 3x in liquid-nitrogen / 37 °C and centrifuged at 14000 g at 4 °C for 5 min. Supernatant was collected in a new tube and stored at -20 °C. Total protein concentration was quantified using the DC assay (Bio-Rad) or Pierce™ BCA Protein Assay Kits (Thermo Scientific) For western blotting, 10 – 20 μg of protein were loaded after dilution in sample buffer 2x. All samples were denatured at 95 °C for 5 min and loaded on polyacrylamide gels (8%, 10%, or 12% depending on the protein to be analysed). For Col1a1 10-20 μg protein was diluted with 2x sample buffer (containing 10% beta-mercaptoethanol) and directly loaded on 10% polyacrylamide gels without heat denaturing (*Iannarone et al., 2019*). The protocol for Western blotting is the same as described previously. Before blocking, the membrane was incubated for 5 min in a Ponceau staining solution (Thermo Scientific) that was used as a loading normalizer. Primary antibodies: Cell Signaling 4858S: p-RPS6 (1: 1000 - 1:2000), Abcam ab2606: 4EBP1 (1: 1000 - 1:2000), Abcam ab27792: p-4EBP1 (1: 1000 - 1:2000), Proteintech 67288-1-Ig: COL1A1 (1: 2000 - 1:5000). For quantification, the total protein stain (TPS) obtained using Ponceau was first determined using ImageJ software (https://imagej.nih.gov/ij/). The protein intensity was normalized to Ponceau signal intensity to compensate for protein loading variation. To determine the relative change in protein expression in Control and SMA samples, the average normalized protein expression for three control samples was calculated. Protein expression was subsequently calculated by dividing the normalized expression in both control and SMA samples by the average normalized expression in control. This returns a normalized, relative expression value for each of the samples analysed. To facilitate reliable comparison of quantifications obtained from different membranes, results were normalized to an internal standard (COL1A1 levels from healthy fibroblast) that was the same across all membranes included in our analyses. The internal standard allows for correction of technical variation caused by transfer, handling, and processing of the individual western blotting membranes. Briefly, Ponceau staining and COL1A1 intensity were first determined using ImageJ. Then, COL1A1 levels were normalized to the Ponceau intensity to control for loading variation. After normalizing COL1A1 expression levels using Ponceau intensity for all samples, including the internal standard. The COL1A1 levels were then divided by the value of the internal standard on each of the membranes, thus allowing comparison of COL1A1 levels across membranes.

### Total RNA extraction from patient’s derived fibroblasts from Coriell Institute of Medical Research

Fibroblasts were grown until reaching 80 % confluency. Each well is washed 3x with cold PBS. Next, 500 µL of Trizol reagent (Invitrogen™) was added to the plates, scraped using a cell scraper, and collected in 1.5 mL tubes. Subsequently, 100 µL of chloroform was added per sample, vortexed, and incubated at room temperature for 5 min. Samples were then centrifuged at 12,000 x g for 15 min. Following centrifugation, the mixture separated into a lower red, phenol-chloroform phase (containing proteins), interphase (containing DNA), and a colourless upper aqueous phase, where RNA exclusively remains.

The upper aqueous phase, containing RNA, was carefully transferred to a fresh tube without disturbing the interphase. For RNA precipitation, 250 µL of isopropyl alcohol and 1 µL of glycoblue (Invitrogen™) were added, mixed by vortexing, and incubated at room temperature for 10 min. Samples were then centrifuged at 12,000 x g for 10 min at 4°C. The supernatant was discarded, and the RNA pellet was washed with 75% ethanol before being centrifuged at 7,500 x g for 10 min at 4°C. Ethanol was removed, and RNA pellets were air-dried and suspended in RNase-free water.

The 500 ng total RNA was treated with DNase I and reverse-transcribed using RNA using RevertAid First Strand cDNA synthesis kit (Thermo Fisher Scientific). The RT-qPCR was performed with 1 μL cDNA for each sample. qPCR was performed with qPCRBIO SyGreen Mix separate-ROX (PCRBiosystem) using the CFX Connect Real-Time PCR Detection System (BioRad) using the primers (**Supplementary Table 5**). qPCR conditions were the same as described previously. The differential expression analysis on total mRNA has been performed by using *FBX038* (*Ramos et al., 2019*) as a normalizer and the ΔΔCT method, as follows:

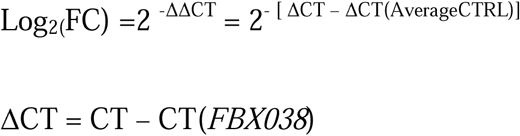

### Ribosome profiling

Tissues from 3 P10 SMA mice and 3 littermates controls were lysed in lysis buffer: 10 mM Tris-HCl pH 7.5, 10 mM MgCl_2_, 10 mM NaCl, 1% w/V TritonX-100, 0.005 U/μL DNAseI, 0.85 U/μL RiboLock RNAse Inhibitor, 1 mM DTT, 200 μg/mL cycloheximide, and 1% w/V Na-deoxycholate. Cytoplasmic lysates were prepared. Lysates were kept for 17-20 min on ice to ensure cell lysis. Then they were centrifuged twice at 12,000 rpm in Eppendorf F-45-30-11 Rotor and Eppendorf Centrifuge 5417R for 10 min and the supernatant was collected. After measuring the absorbance at 260 nm, the concentration of NaCl was adjusted to a final concentration of 100 mM. Lysates were treated using RNase I (10U per unit of absorbance at 260nm in the lysate) at room temperature for 45 min. The reaction was stopped by adding 100 U of SUPERase-In RNase inhibitor (Thermo Fisher Scientific) and lysates were loaded on 13.2 mL polyallomer ultracentrifuge tubes (Beckman) containing 10%-40% (w/V) linear sucrose gradient. The samples were ultracentrifuged for 1 h 30 min at 40,000 rpm at 4°C in Beckman Optima XPN-100 Ultracentrifuge in SW41 rotor. Using a fractionator, the fraction containing monosomes with ribosome-protected fragments (RPFs) was isolated and RNA was purified from the fraction using the phenol/chloroform protocol as described in *Lauria et al., 2020*. RPFs of 28-32 nt in length were obtained by excision of the corresponding band on a denaturing 15% Urea-TBE gel. The excised gel was transferred into a 1.5ml tube for RNA extraction. DEPC water (400 µL), supplemented with 40 µL of 5M CH_3_COONH_4,_ and 2 µL of 10% SDS were added to the gel and then smashed using sterile pestles. The fragmented gel was incubated under rotation on a wheel overnight at 4 °C. The following day samples were transferred into the filter tube and centrifuged for 3 min at 2,300 g to separate any gel residue. RPF was extracted from the solution using the phenol/chloroform method as described in *Lauria et al., 2020*. The RPFs were dephosphorylated using T4 polynucleotide kinase (New England Biolabs #M0201S) for 1 hr at 37 °C. These RPFs were used to prepare the Ribo-seq Library using Takara’s small library preparation kit (SMARTer smRNA-seq kit) following manufacturer instructions. The resulting libraries were sequenced using Illumina NOVA-Seq 6000 at the CIBIO NGS facility of the University of Trento, Italy. Experiments were performed in triplicate.

### Preprocessing of ribosome profiling data

The 3’ adapters (CTGTAGGCACCATCAAT) and overrepresented sequences (reverse complement of “Illumina sequencing primers”: CTCTTTCCCTACACG, ACGCTCTTCCGATCT) were clipped. Three occurrences of the adapter sequence are searched. The first 5’ nucleotide was trimmed. Reads shorter than 19 nt were also discarded. Maximum error rate and overlap parameters were set at 0.15 and 10, respectively. Reads mapping on the collection of *Mus musculus* rRNAs (from the SILVA rRNA database, release 119) and tRNAs (from the Genomic tRNA database: gtrnadb.ucsc.edu/) were removed. The remaining reads were mapped on the mouse genome (using the Gencode M22 annotations - ensembl 97) allowing a maximum of 5 multiple alignments for each read. All alignments were performed with STAR (v2.5.3a) using default settings.

Reads mapping on the same position with respect to transcript coordinates were then removed before proceeding with positional analyses. Ribosome profiling analyses based on positional information were performed using the R package riboWaltz (*Lauria et al., 2018*). Duplicated reads mapping on the same position with respect to transcript coordinates were removed.

First, the genes with zero counts for all the replicates were not considered for further analysis. Next, *fpkm* was computed (with the *rpkm* function provided within the EdgeR package) (*Chen et al., 2016*; *Lun et al., 2016*; *McCarthy et al., 2012*; *Robinson et al., 2010*) using as gene length parameter the length of each gene retrieved from the GTF file used during the alignment step. The value corresponding to the 80^th^ quantile of the *fpkm* distribution for each replicate was used as a cut-off in the filtering step. Genes with *fpkm* values above the threshold values for all the replicates of a condition (control or SMA) were kept. Finally, genes with a signal lower than a replicate-specific threshold were filtered out.

Normalization among replicates was performed with the trimmed mean of M-values normalization method (TMM) implemented in the EdgeR Bioconductor package. Pairwise differential analyses (SMA vs CTRL) were performed independently for each. Genes showing significantly differential ribosome occupancy were defined by the following cut-off values: *cpm threshold*=0.05, *|log2FC|>0.3*, *p-value <*0.05.

### Enrichment Analysis

Gene ontology and KEGG analysis was computed on the set of genes with increased and decreased ribosome occupancy using the Enrichr resource with default settings (http://amp.pharm.mssm.edu/Enrichr/).

### Network and Communities Analysis

Protein-protein interaction analyses were performed using STRING (https://string-db.org/).

### Statistical analysis

All plots represent mean ± standard error of the mean. All data were tested for normality using the Shapiro-Wilk test. Two-tailed Student’s t-tests (unpaired) were performed for pairwise comparisons of control and SMA groups. For multiple comparisons between healthy and SMA fibroblast lines, a one-way ANOVA test with Holm correction of p-values was performed. For treatment outcomes, a two-tailed two-sample Student’s t-test (unpaired) with Holm’s correction of p-values was performed. Significance was set at p ≤ 0.05 for *, p ≤ 0.01 for **, p ≤ 0.001 for *** and p ≤ 0.0001 for ****. For all experiments, quantification and statistics were derived from n≥3 independent replicates unless specified in the legends.

## Data Availability

All data sets for ribosome profiling sequencing experiment can be provided by the corresponding author after request.

## Author Contributions

GV and THG designed the project. GV, EG, PC and THG obtained funding. GV, THG and EG planned the experiments. GS, EP, MP, YTH, CP, IS, FM, DD, HC, EG, MB, FL performed the experiments, analysed the data and organised the figures. THG, MB and RK provided the tissues and valuable discussions. GS and GV drafted the manuscript and all authors contributed to writing. All authors have read and agreed to the published version of the manuscript.

## Acknowledgement

We are grateful to Dr. Lyndsay Murray for useful discussions. We would like to thank the staff at the Core Facilities Next Generation Sequencing Facility (NGS) at Department CIBIO University of Trento, Italy; and Michela Laferla, Francesca Spanò, Elena Gerola and Rosaanna Cascone of the Administrative staff at IBF-CNR for crucial support with administrative management, and Marta Marchioretto for her valuable help as Lab Manager at IBF-CNR.

## Funding

GV, EG, LvDP, THG and PC are grateful for funding from the European Union’s Horizon 2020 Research and Innovation Program under the Marie Sklodowska-Curie grant (H2020 Marie Sklodowska-Curie Actions) agreement no. 956185 (SMABEYOND ITN). GV is grateful for funding from AFM Telethon (#23692), Telethon (GGP19115 and GMR23T1048), EU funding within the MUR PNRR ‘National Center for Gene Therapy and Drugs based on RNA Technology’ (Project no. CN00000041 CN3 RNA) and Caritro Foundation (Rif.Int. 2021.0571).

## Conflict of interest

The authors declare no conflict of interest.

## SUPPLEMENTARY MATERIAL

**Supplementary Figure 1.**
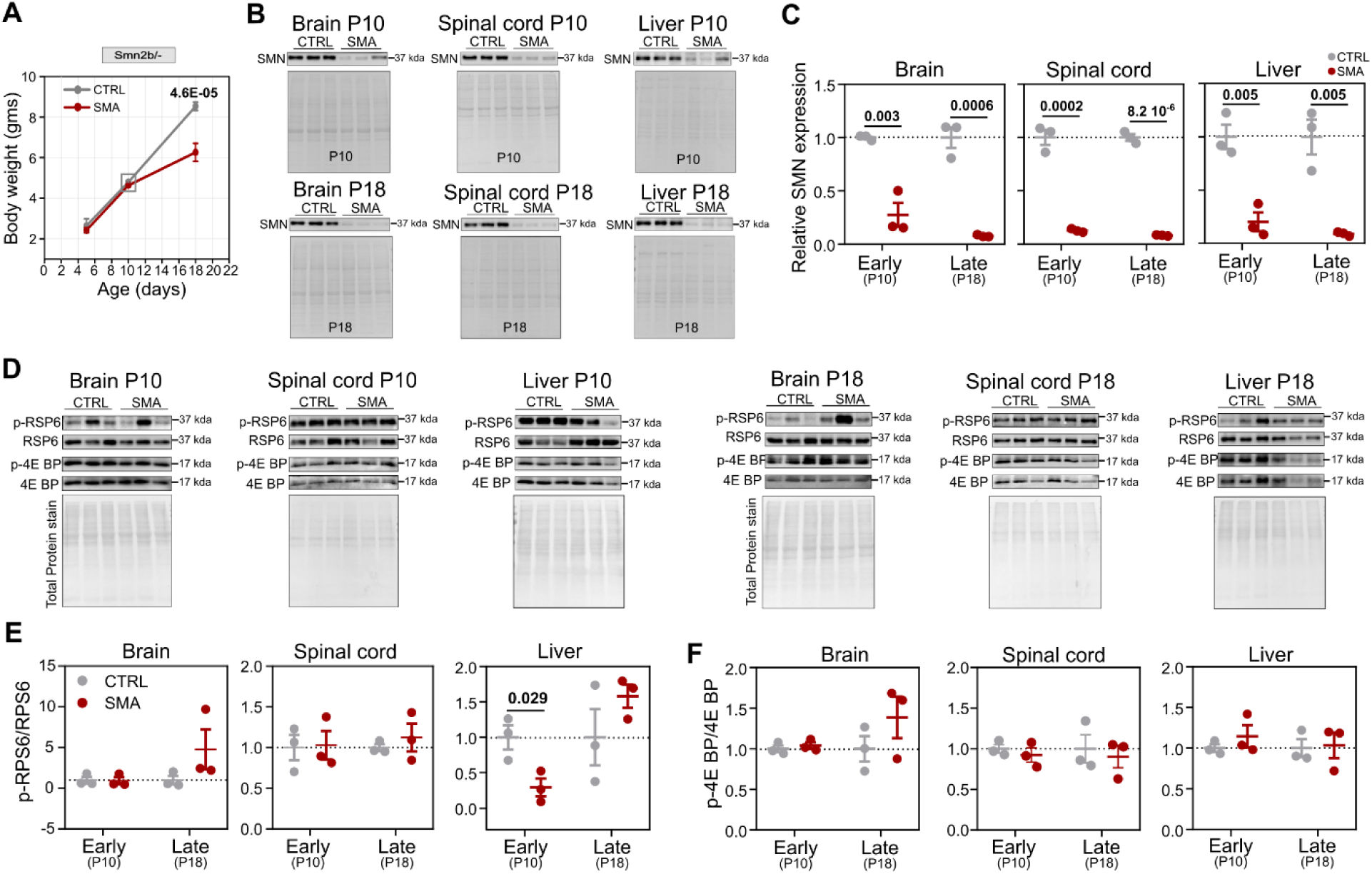
**(A)** Comparison of the body weight between control and Smn^2B/-^ mice. The grey bo represents the pre-symptomatic stage of SMA. Data are meanD±Ds.e.m. of nD≥D3 biologically independent experiments. Significance was calculated by unpaired two-tailed Student’s t-test. Significant p-values are reporte on the plots. **(B)** Comparison of SMN protein expression between control and SMA tissues using western blot of the total protein lysate from the brain, spinal cord, and liver at P10 and P18 from Smn^2B/-^ mice. **(C)** Quantification of SMN protein levels relative to total protein stain and to control conditions in the brain, spinal cord, and liver. SMN expression values were normalized and compared with control values for each tissue. Data are meanD±Ds.e.m. of nD=D3 biologically independent experiments. Significance was calculated by unpaired two-tailed Student’s t-test. Significant p-values are reported on the plots. **(D)** Western blot analysis for p-RPS6, RPS6, p-4EBP1, and 4EBP1 proteins in brain, spinal cord, and liver of the Control and SMA at P10 and P18 stages. **(E,F)** Quantification of western blot from control (grey) and SMA (red) for the p-RPS6/RPS6 (E) and p-4EBP1/4EBP1 (F). Data are plotte as mean ± s.e.m. of n = 3 mice. Significant differences were assessed using an unpaired two-tailed Student’s t-test. Significant p-values are reported on the plots.

**Supplementary Figure 2.**
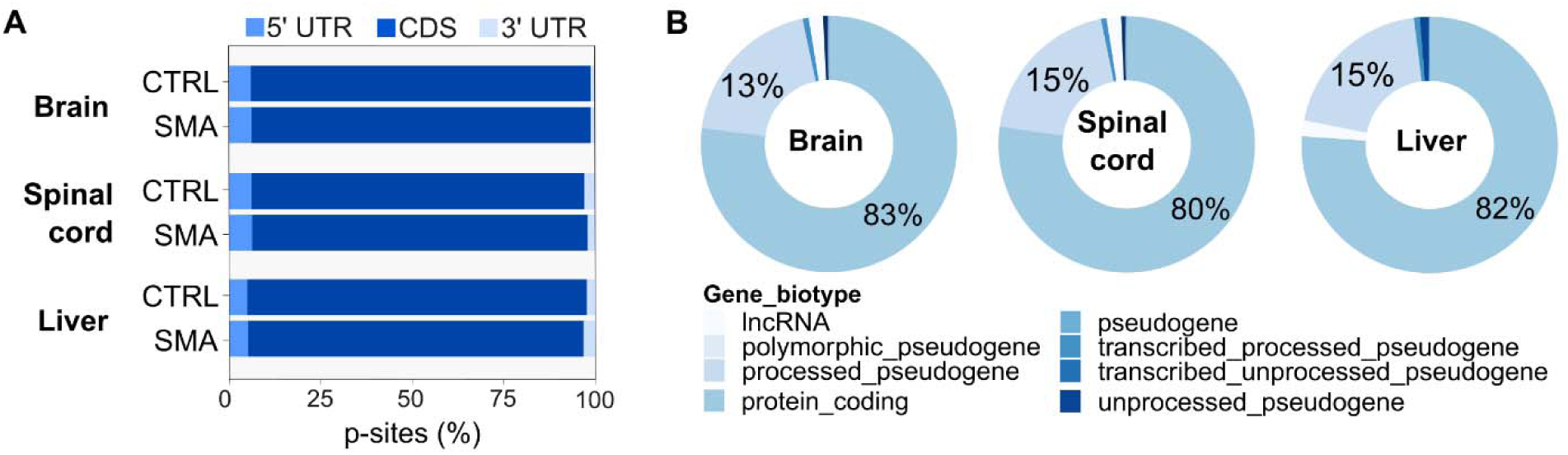
**(A)** Positional enrichment along the three mRNA regions of the CDS. Data are shown as the percentage of p-sites in the 5′ UTR, CDS, and 3′ UTR, averaged across the replicates. **(B)** Transcript types enriched in fragments protected by ribosomes are predominantly associated with protein-coding genes.

**Supplementary Figure 3.**
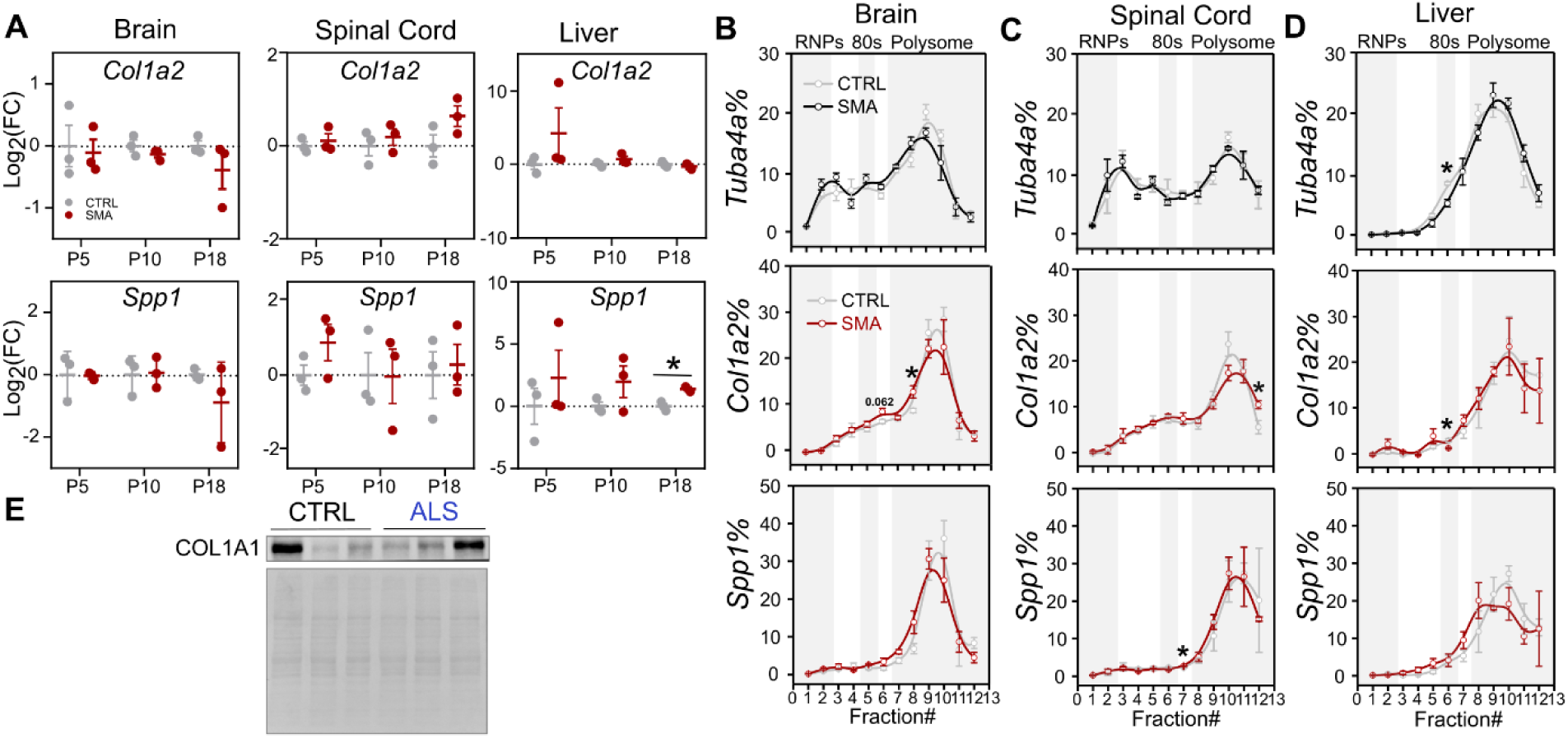
**(A)** Quantification of qPCR analysis performed for the *Col1a2*, and *Spp1* transcripts in cytoplasmic mRNAs from control and SMA brain, spinal cord, and liver at the P5, P10, and P18 stages of SMA disease in Smn^2B/-^ mice. Data are mean ± s.e.m. among n = 3 biologically independent samples. Significant changes were assessed using unpaired two-sided t-tests. * p-val<0.05. **(B-D)** Relative co-sedimentation profile of *Tuba4a, Col1a2*, and *Spp1* mRNAs along the sucrose gradient fractions of control and SMA brain (**B**), spinal cord (**C**), an liver (**D**) at P10. Data are mean ± s.e.m. among n = 3 biologically independent samples. Significant changes were assessed using unpaired two-sided Student’s t-tests. * p-val<0.05. **(E)** Protein levels of COL1A1 in the cortex and spinal cord of 3-month-old ALS and littermate controls mice using Western Blot.

**Supplementary Figure 4.**
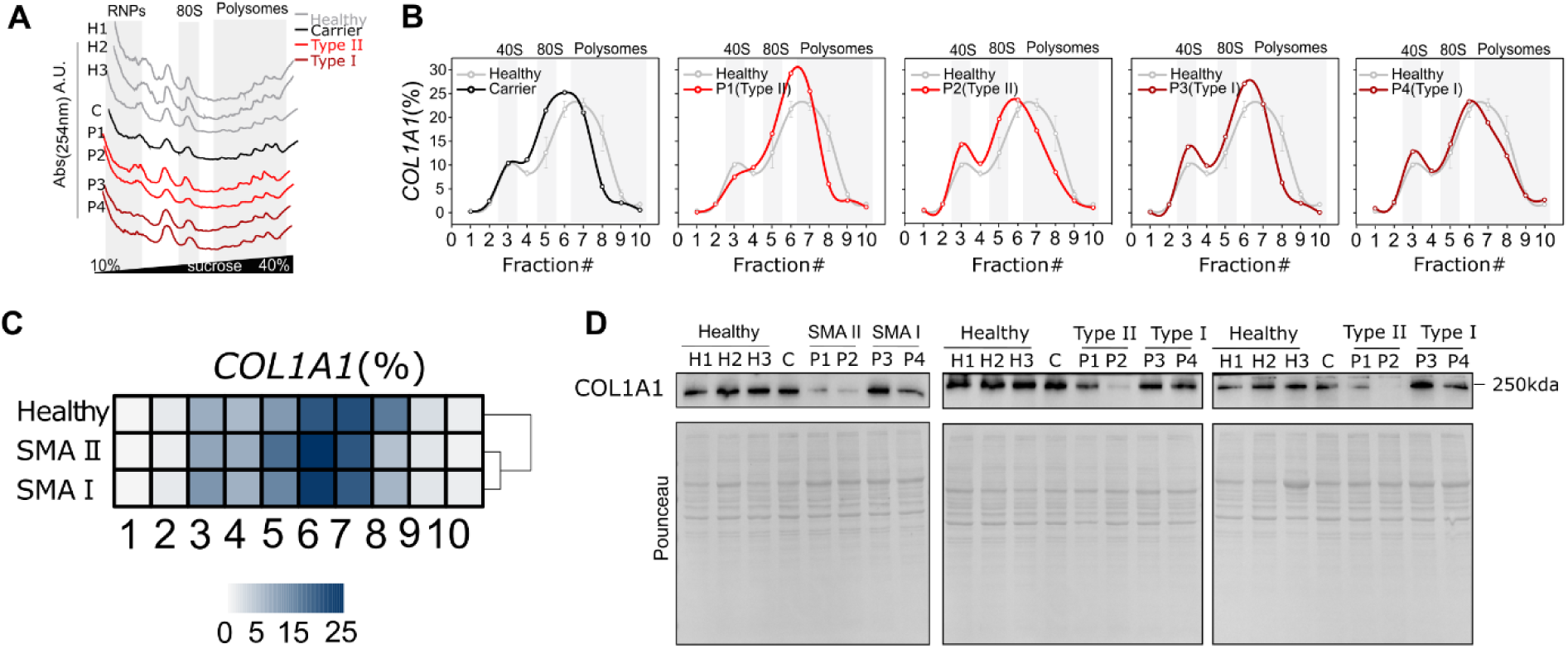
**(A)** Polysome profiles of healthy, carrier, and SMA patient-derived fibroblasts. **(B)** Heat map representing the co-sedimentation profile of *COL1A1* mRNA transcript in healthy controls and SMA patient-derived fibroblasts. **(C)** Comparison between the relative co-sedimentation profile of *COL1A1* mRNA from healthy controls (n=3), carrier, and individual SMA patient-derived fibroblasts. **(D)** COL1A1 protein immunoblot i healthy, carrier, and individual SMA patient-derived fibroblasts.

**Supplementary Figure 5.**
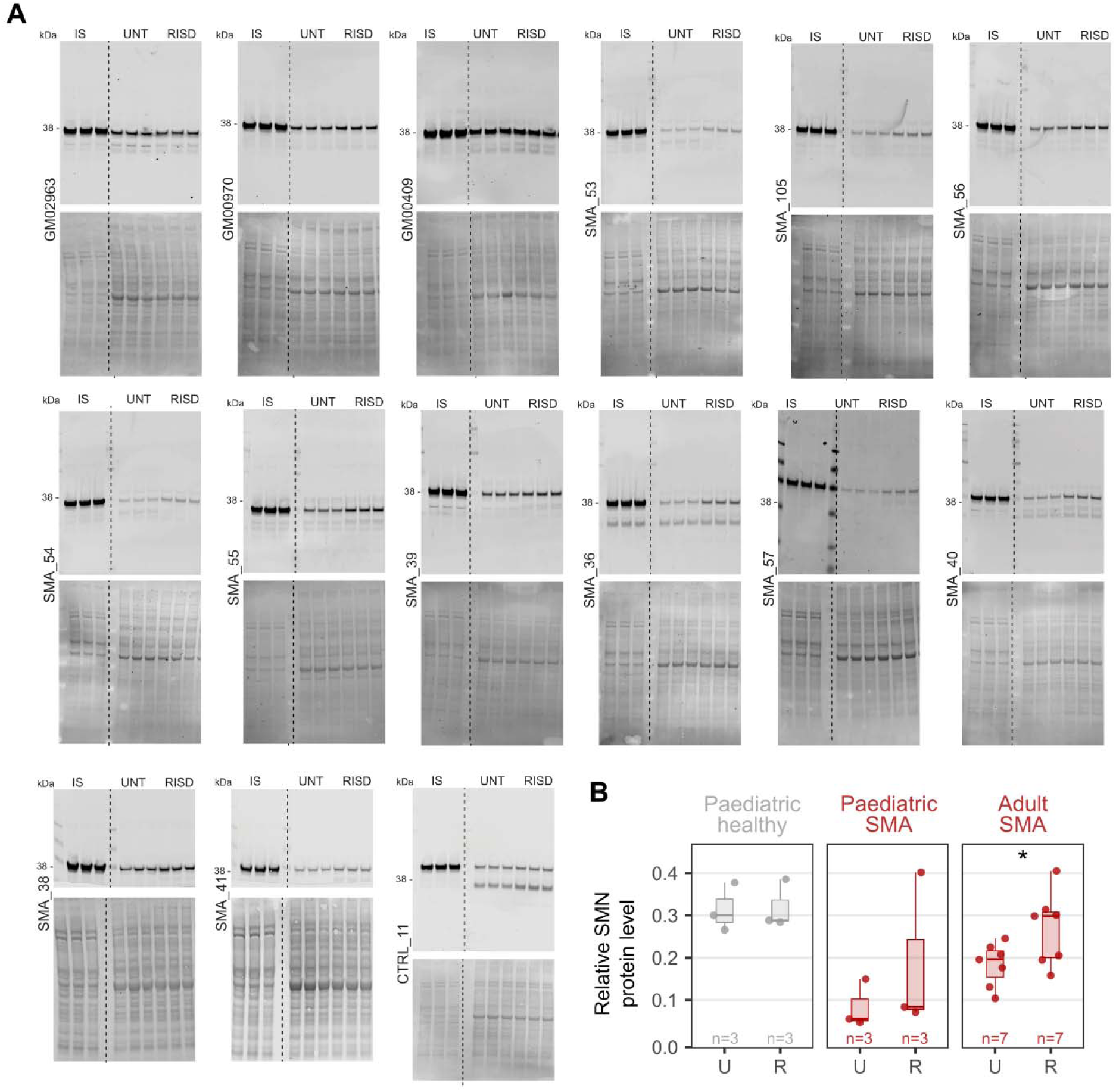
**(A)** Representative western blot images showing SMN protein and total protein lysates from control (CTRL) and SMA patient-derived fibroblast samples. Samples include untreated (UNT) and risdiplam-treated (RISD) conditions. Internal standard (IS) is used to compare the SMN across the different membranes. Each blot is accompanied by a corresponding total protein stain (lower panel). Sample labels above each lane denote the specific fibroblast line used in the analysis (See Supplementary Table 4). (**B**) Comparison of SMN protein expression from control (CTRL) and SMA patient-derived fibroblast samples. Data represent the mean ± s.e.m. of n individual, shown as a dot, for each group. Significant changes were assessed using paired two-sided t-tests, with Holm correction of p-values.

**Supplementary Figure 6.**
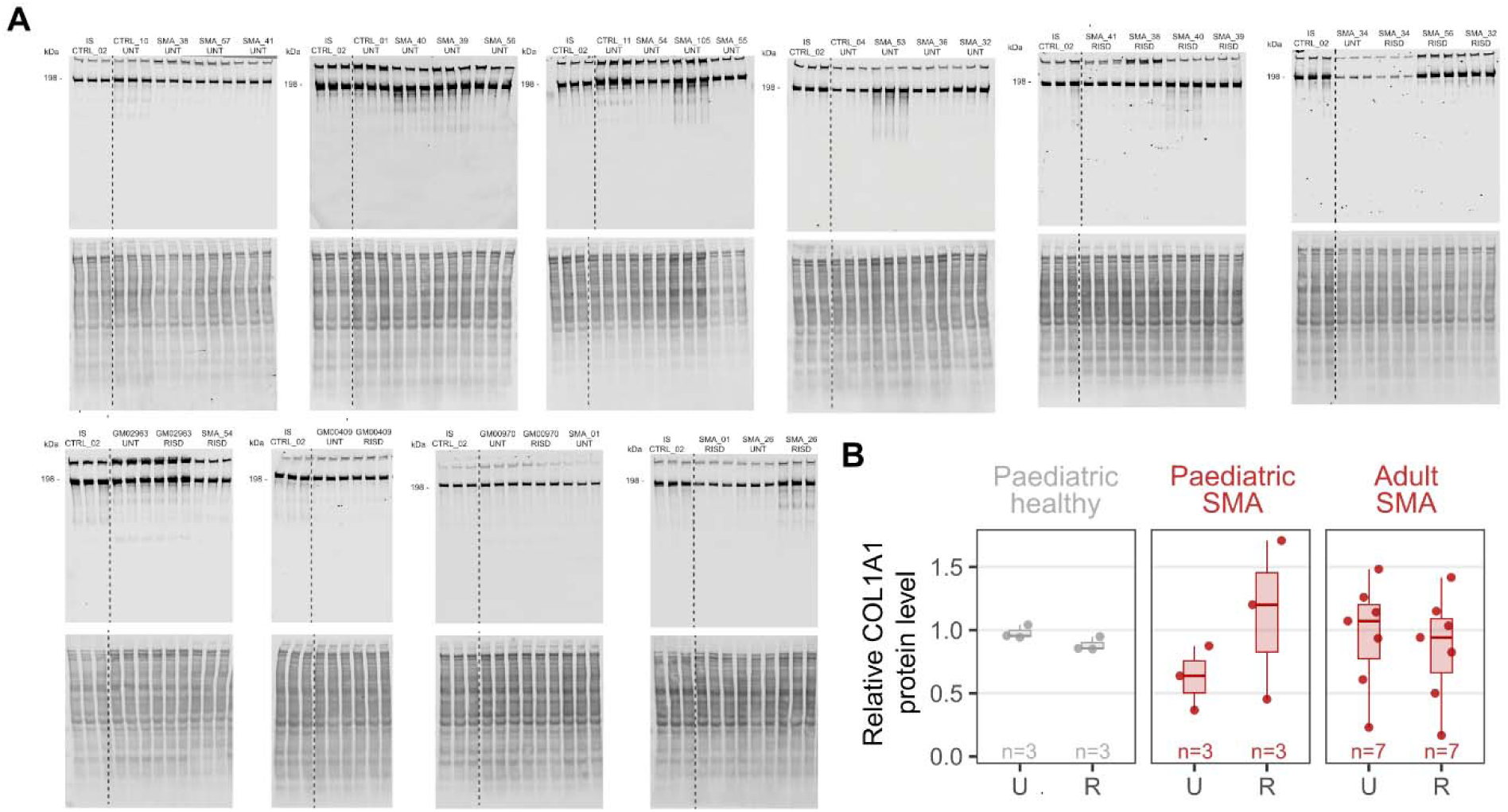
(**A**) Representative western blot images showing COL1A1 protein and total protei lysates from control (CTRL) and SMA patient-derived fibroblast samples. Samples include untreated (UNT) an risdiplam-treated (RISD) conditions. Internal standard (IS) is used to compare the COL1A1 across the different membranes. Each blot is accompanied by a corresponding total protein stain (lower panel). Sample labels abov each lane denote the specific fibroblast line used in the analysis (See Supplementary Table 4). (**B**) Comparison of COL1A1 protein expression from control (CTRL) and SMA patient-derived fibroblast samples. Data represent the mean ± s.e.m. of n individual, shown as a dot, for each group. Significant changes were assessed using paired two-sided t-tests, with Holm correction of p-values.

**Supplementary Table 1:**
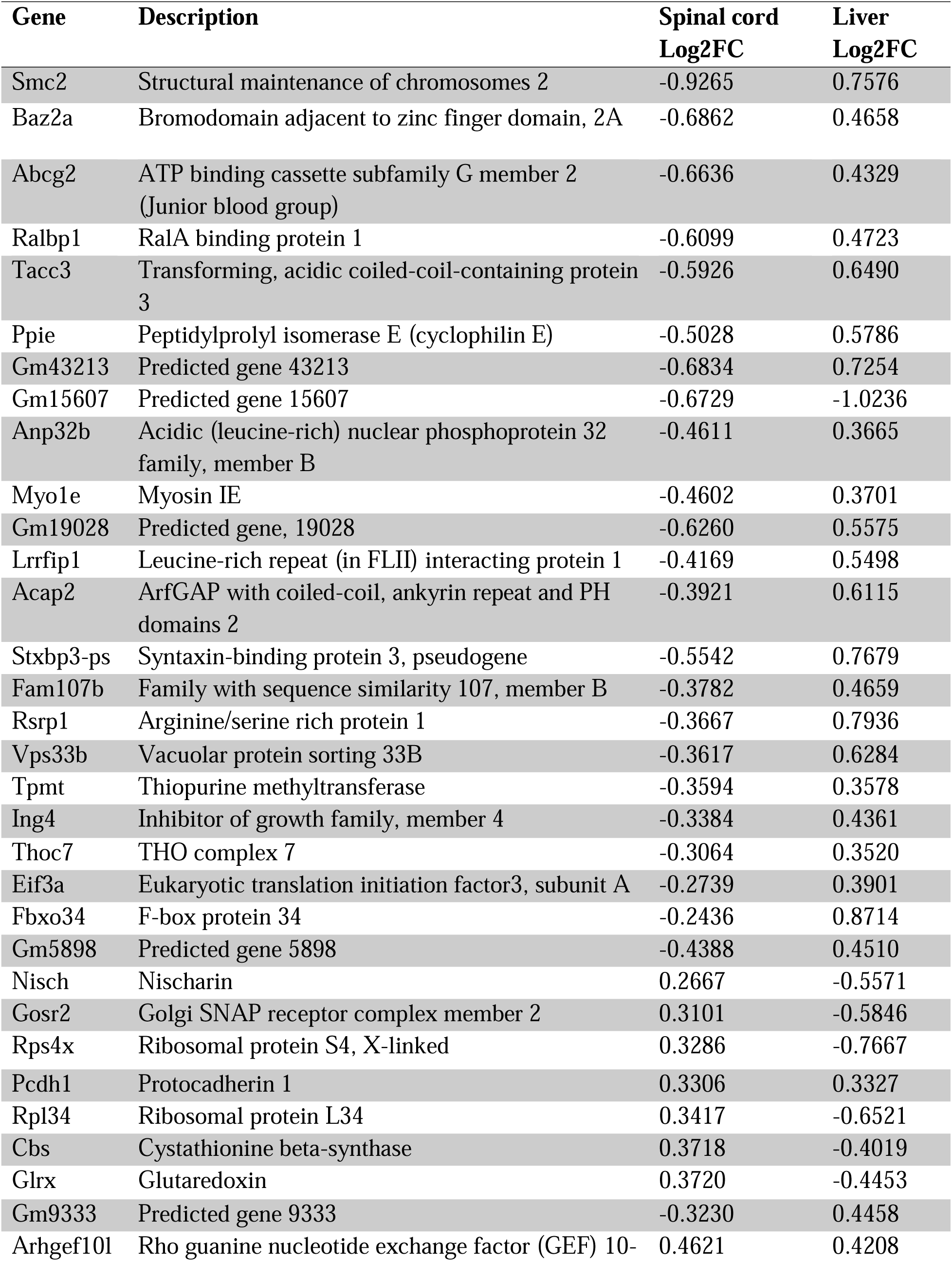
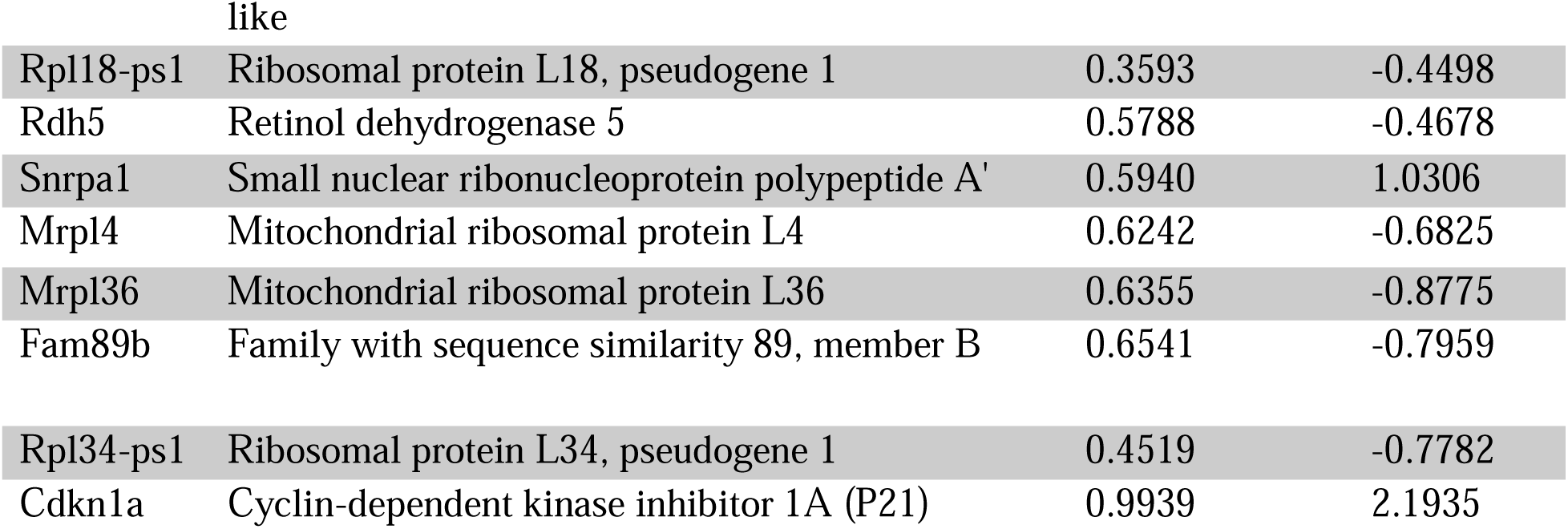
List of shared transcripts between Spinal cord and Liver.

| **Gene** | **Description** | **Spinal cord**<br>**Log2FC** | **Liver**<br>**Log2FC** |
| --- | --- | --- | --- |
| Smc2 | Structural maintenance of chromosomes 2 | -0.9265 | 0.7576 |
| Baz2a | Bromodomain adjacent to zinc finger domain, 2A | -0.6862 | 0.4658 |
| Abcg2 | ATP binding cassette subfamily G member 2<br>(Junior blood group) | -0.6636 | 0.4329 |
| Ralbp1 | RalA binding protein 1 | -0.6099 | 0.4723 |
| Tacc3 | Transforming, acidic coiled-coil-containing protein 3 | -0.5926 | 0.6490 |
| Ppie | Peptidylprolyl isomerase E (cyclophilin E) | -0.5028 | 0.5786 |
| Gm43213 | Predicted gene 43213 | -0.6834 | 0.7254 |
| Gm15607 | Predicted gene 15607 | -0.6729 | -1.0236 |
| Anp32b | Acidic (leucine-rich) nuclear phosphoprotein 32 family, member B | -0.4611 | 0.3665 |
| Myo1e | Myosin IE | -0.4602 | 0.3701 |
| Gm19028 | Predicted gene, 19028 | -0.6260 | 0.5575 |
| Lrrfip1 | Leucine-rich repeat (in FLII) interacting protein 1 | -0.4169 | 0.5498 |
| Acap2 | ArfGAP with coiled-coil, ankyrin repeat and PH domains 2 | -0.3921 | 0.6115 |
| Stxbp3-ps | Syntaxin-binding protein 3, pseudogene | -0.5542 | 0.7679 |
| Fam107b | Family with sequence similarity 107, member B | -0.3782 | 0.4659 |
| Rsrp1 | Arginine/serine rich protein 1 | -0.3667 | 0.7936 |
| Vps33b | Vacuolar protein sorting 33B | -0.3617 | 0.6284 |
| Tpmt | Thiopurine methyltransferase | -0.3594 | 0.3578 |
| Ing4 | Inhibitor of growth family, member 4 | -0.3384 | 0.4361 |
| Thoc7 | THO complex 7 | -0.3064 | 0.3520 |
| Eif3a | Eukaryotic translation initiation factor3, subunit A | -0.2739 | 0.3901 |
| Fbxo34 | F-box protein 34 | -0.2436 | 0.8714 |
| Gm5898 | Predicted gene 5898 | -0.4388 | 0.4510 |
| Nisch | Nischarin | 0.2667 | -0.5571 |
| Gosr2 | Golgi SNAP receptor complex member 2 | 0.3101 | -0.5846 |
| Rps4x | Ribosomal protein S4, X-linked | 0.3286 | -0.7667 |
| Pcdh1 | Protocadherin 1 | 0.3306 | 0.3327 |
| Rpl34 | Ribosomal protein L34 | 0.3417 | -0.6521 |
| Cbs | Cystathionine beta-synthase | 0.3718 | -0.4019 |
| Glrx | Glutaredoxin | 0.3720 | -0.4453 |
| Gm9333 | Predicted gene 9333 | -0.3230 | 0.4458 |
| Arhgef10l | Rho guanine nucleotide exchange factor (GEF) 10- | 0.4621 | 0.4208 |
| Rpl18-ps1 | Ribosomal protein L18, pseudogene 1 | 0.3593 | -0.4498 |
| Rdh5 | Retinol dehydrogenase 5 | 0.5788 | -0.4678 |
| Snrpa1 | Small nuclear ribonucleoprotein polypeptide A' | 0.5940 | 1.0306 |
| Mrpl4 | Mitochondrial ribosomal protein L4 | 0.6242 | -0.6825 |
| Mrpl36 | Mitochondrial ribosomal protein L36 | 0.6355 | -0.8775 |
| Fam89b | Family with sequence similarity 89, member B | 0.6541 | -0.7959 |
| Rpl34-ps1 | Ribosomal protein L34, pseudogene 1 | 0.4519 | -0.7782 |
| Cdkn1a | Cyclin-dependent kinase inhibitor 1A (P21) | 0.9939 | 2.1935 |

**Supplementary Table 2:** List of shared transcripts between Brain and Spinal cord.

| **Gene** | **Description** | **Brain**<br>**LogFC** | **Spinal cord**<br>**LogFC** |
| --- | --- | --- | --- |
| Smn1 | Survival motor neuron 1 | -1.4189 | -1.0416 |
| Meig1 | Meiosis expressed gene 1 | 0.3937 | -0.6623 |
| Mettl17 | Methyltransferase like 17 | 0.6503 | -0.4972 |
| Spp1 | Secreted phosphoprotein 1 | -1.0508 | -0.3213 |
| Ninj1 | Ninjurin 1 | -0.3787 | 0.4761 |
| Tmub1 | Transmembrane and ubiquitin-like domain<br>containing 1 | -0.6031 | 0.6976 |

**Supplementary Table 3:**
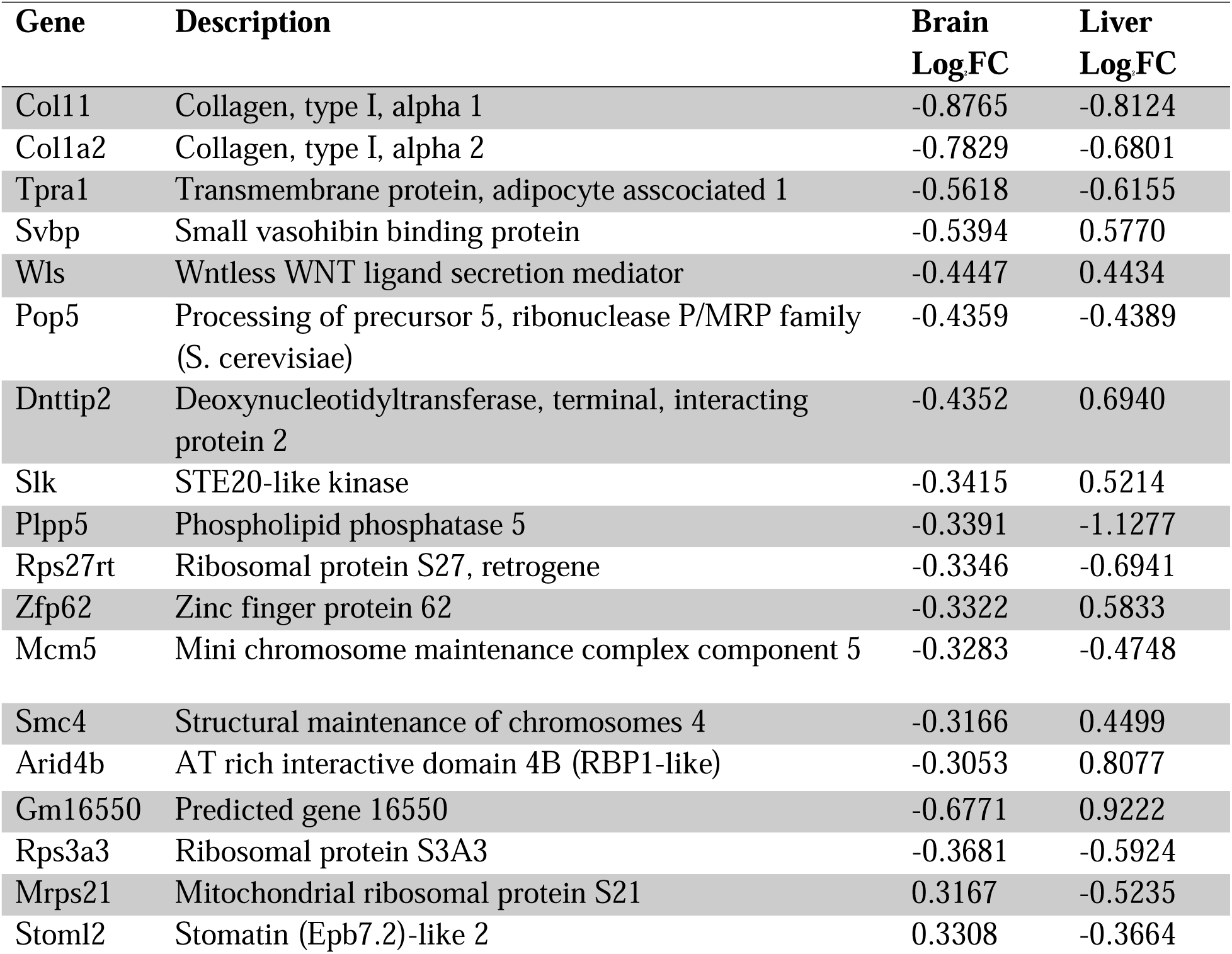
List of shared transcripts between Brain and Liver.

**Supplementary Table 4:**
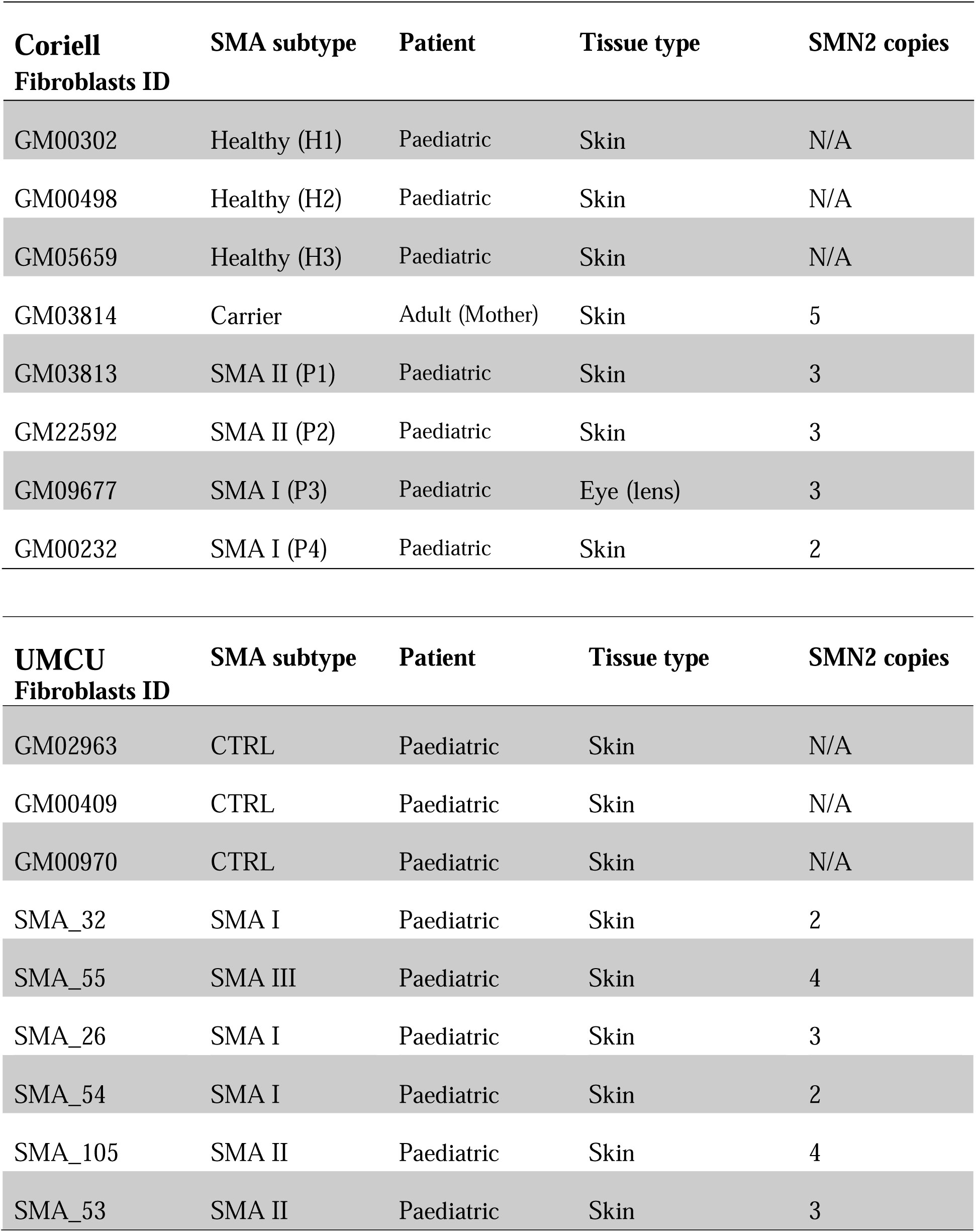
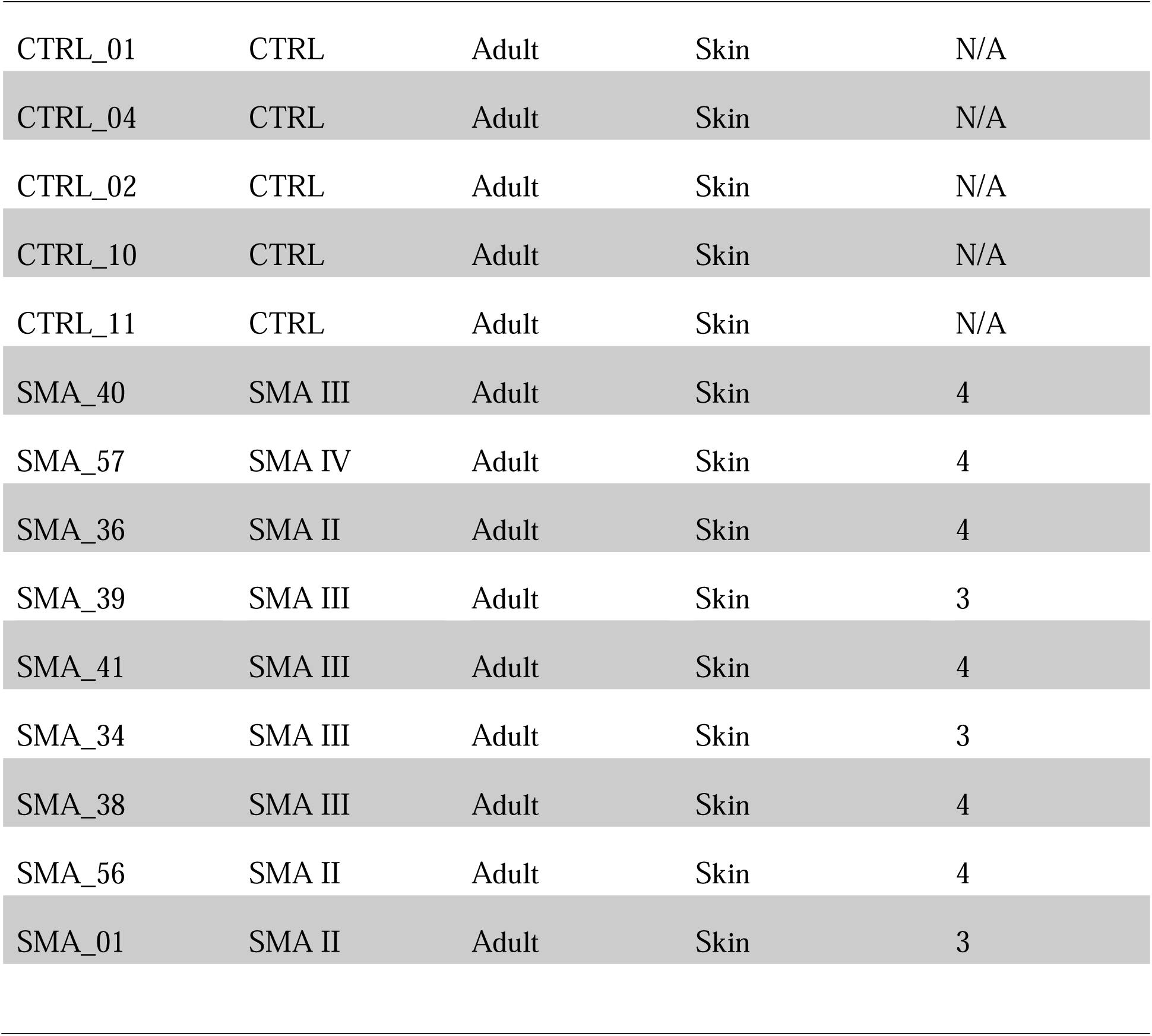
Characteristics of SMA Patient Fibroblasts cohort from Coriell Institute of Medical Research and University Medical Center Utrecht (UMCU)

| **Coriell**<br>**Fibroblasts ID** | **SMA subtype** | **Patient** | **Tissue type** | **SMN2 copies** |
| --- | --- | --- | --- | --- |
| GM00302 | Healthy (H1) | Paediatric | Skin | N/A |
| GM00498 | Healthy (H2) | Paediatric | Skin | N/A |
| GM05659 | Healthy (H3) | Paediatric | Skin | N/A |
| GM03814 | Carrier | Adult (Mother) | Skin | 5 |
| GM03813 | SMA II (P1) | Paediatric | Skin | 3 |
| GM22592 | SMA II (P2) | Paediatric | Skin | 3 |
| GM09677 | SMA I (P3) | Paediatric | Eye (lens) | 3 |
| GM00232 | SMA I (P4) | Paediatric | Skin | 2 |
| **UMCU**<br>**Fibroblasts ID** | **SMA subtype** | **Patient** | **Tissue type** | **SMN2 copies** |
| GM02963 | CTRL | Paediatric | Skin | N/A |
| GM00409 | CTRL | Paediatric | Skin | N/A |
| GM00970 | CTRL | Paediatric | Skin | N/A |
| SMA_32 | SMA I | Paediatric | Skin | 2 |
| SMA_55 | SMA III | Paediatric | Skin | 4 |
| SMA_26 | SMA I | Paediatric | Skin | 3 |
| SMA_54 | SMA I | Paediatric | Skin | 2 |
| SMA_105 | SMA II | Paediatric | Skin | 4 |
| SMA_53 | SMA II | Paediatric | Skin | 3 |
| CTRL_01 | CTRL | Adult | Skin | N/A |
| CTRL_04 | CTRL | Adult | Skin | N/A |
| CTRL_02 | CTRL | Adult | Skin | N/A |
| CTRL_10 | CTRL | Adult | Skin | N/A |
| CTRL_11 | CTRL | Adult | Skin | N/A |
| SMA_40 | SMA III | Adult | Skin | 4 |
| SMA_57 | SMA IV | Adult | Skin | 4 |
| SMA_36 | SMA II | Adult | Skin | 4 |
| SMA_39 | SMA III | Adult | Skin | 3 |
| SMA_41 | SMA III | Adult | Skin | 4 |
| SMA_34 | SMA III | Adult | Skin | 3 |
| SMA_38 | SMA III | Adult | Skin | 4 |
| SMA_56 | SMA II | Adult | Skin | 4 |
| SMA_01 | SMA II | Adult | Skin | 3 |

**Supplementary Table 5:**
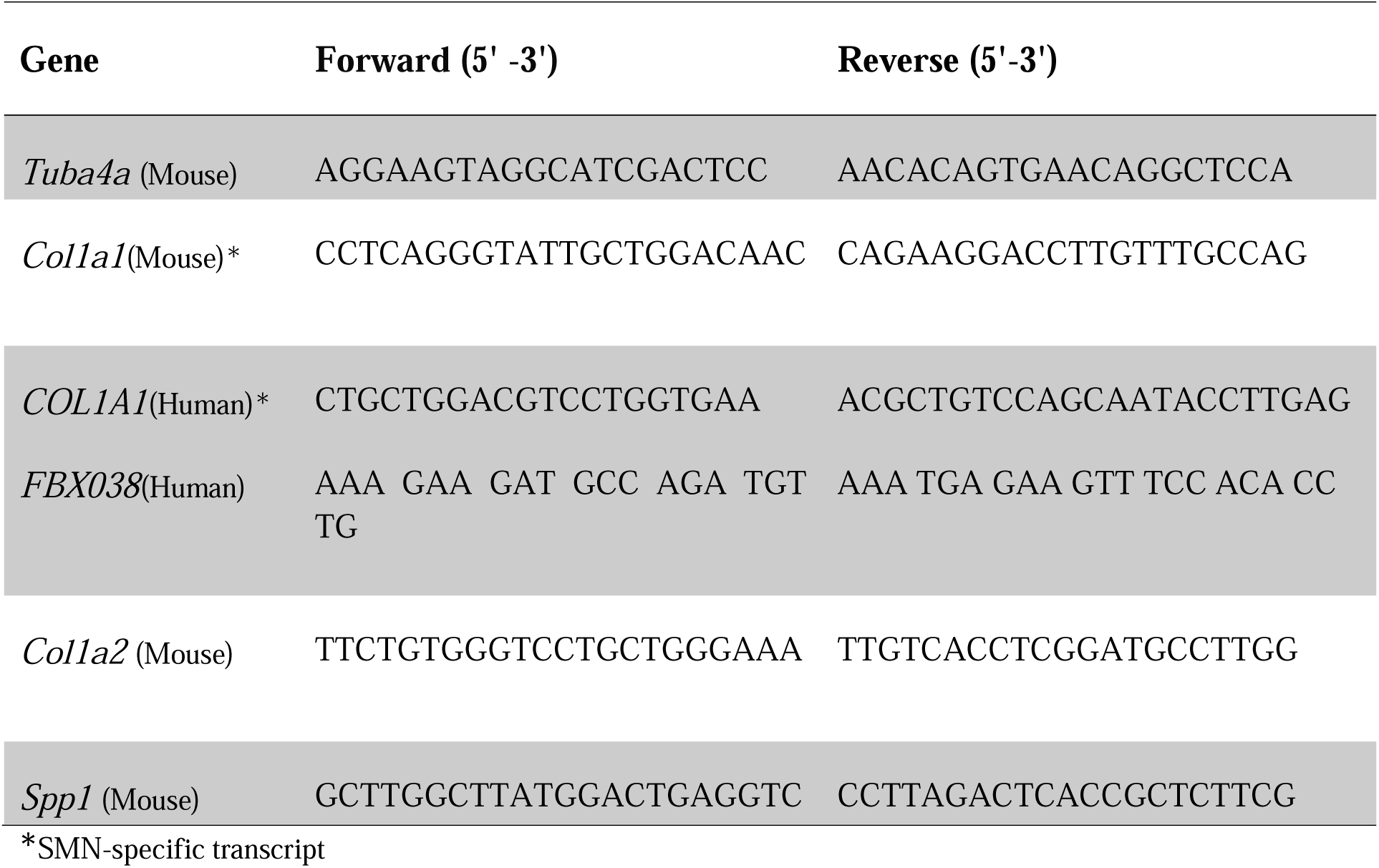
Primer used for RT-qPCR.

## Notes

### Competing Interest Statement

The authors have declared no competing interest.

### Summary of Updates

Correction in the name of a co- author and ORCID.

## REFERENCES

Allardyce H, Kuhn D, Hernandez-Gerez E, Hensel N, Huang YT, Faller K, Gillingwater TH, Quondamatteo F, Claus P, Parson SH. Renal pathology in a mouse model of severe Spinal Muscular Atrophy is associated with downregulation of Glial Cell-Line Derived Neurotrophic Factor (GDNF). Hum Mol Genet. 2020 Aug 11;29(14):2365–2378.

Almeida MMA, De Repentigny Y, Gagnon S, Sutton ER, Kothary R. Impact of liver-specific survival motor neuron (SMN) depletion on central nervous system and peripheral tissue pathology. Elife. 2025 Feb 20;13:RP99141.

Arnold ES, Ling SC, Huelga SC, Lagier-Tourenne C, Polymenidou M, Ditsworth D, Kordasiewicz HB, McAlonis-Downes M, Platoshyn O, Parone PA, Da Cruz S, Clutario KM, Swing D, Tessarollo L, Marsala M, Shaw CE, Yeo GW, Cleveland DW. ALS-linked TDP-43 mutations produce aberrant RNA splicing and adult-onset motor neuron disease without aggregation or loss of nuclear TDP-43. Proc Natl Acad Sci U S A. 2013 Feb 19;110(8):E736–45. doi: 10.1073/pnas.1222809110.

Baranello G, Vai S, Broggi F, Masson R, Arnoldi MT, Zanin R, Mastella C, Bianchi ML. Evolution of bone mineral density, bone metabolism and fragility fractures in Spinal Muscular Atrophy (SMA) types 2 and 3. Neuromuscul Disord. 2019 Jul;29(7):525–532.

Baranello G, Gorni K, Daigl M, Kotzeva A, Evans R, Hawkins N, Scott DA, Mahajan A, Muntoni F, Servais L. Prognostic Factors and Treatment-Effect Modifiers in Spinal Muscular Atrophy. Clin Pharmacol Ther. 2021 Dec;110(6):1435–1454.

Baranello G; Neurodevelopment in SMA Working Group. The emerging spectrum of neurodevelopmental comorbidities in early-onset Spinal Muscular Atrophy. Eur J Paediatr Neurol. 2024 Jan;48:67–68

Bayoumy S, Verberk IMW, Vermunt L, Willemse E, den Dulk B, van der Ploeg AT, Pajkrt D, Nitz E, van den Hout JMP, van der Post J, Wolf NI, Beerepoot S, Groen EJN, Tüngler V, Teunissen CE. Neurofilament light protein as a biomarker for spinal muscular atrophy: a review and reference ranges. Clin Chem Lab Med. 2024 Jan 15;62(7):1252–1265.

Bäumer D, Lee S, Nicholson G, Davies JL, Parkinson NJ, Murray LM, Gillingwater TH, Ansorge O, Davies KE, Talbot K. Alternative splicing events are a late feature of pathology in a mouse model of spinal muscular atrophy. PLoS Genet. 2009 Dec;5(12):e1000773.

Bernabò P, Tebaldi T, Groen EJN, Lane FM, Perenthaler E, Mattedi F, Newbery HJ, Zhou H, Zuccotti P, Potrich V, Shorrock HK, Muntoni F, Quattrone A, Gillingwater TH, Viero G. In Vivo Translatome Profiling in Spinal Muscular Atrophy Reveals a Role for SMN Protein in Ribosome Biology. Cell Rep. 2017 Oct 24;21(4):953–965.

Bowerman M, Murray LM, Beauvais A, Pinheiro B, Kothary R. A critical smn threshold in mice dictates onset of an intermediate spinal muscular atrophy phenotype associated with a distinct neuromuscular junction pathology. Neuromuscul Disord. 2012 Mar;22(3):263–76.

Bowerman M, Shafey D, Kothary R. Smn depletion alters profilin II expression and leads to upregulation of the RhoA/ROCK pathway and defects in neuronal integrity. J Mol Neurosci. 2007;32(2):120–31.

Brown SJ, Kline RA, Synowsky SA, Shirran SL, Holt I, Sillence KA, Claus P, Wirth B, Wishart TM, Fuller HR. The Proteome Signatures of Fibroblasts from Patients with Severe, Intermediate and Mild Spinal Muscular Atrophy Show Limited Overlap. Cells. 2022 Aug 23;11(17):2624.

Chaytow H, Faller KME, Huang YT, Gillingwater TH. Spinal muscular atrophy: From approved therapies to future therapeutic targets for personalized medicine. Cell Rep Med. 2021 Jul 21;2(7):100346.

Chen Y, Lun AT, Smyth GK. From reads to genes to pathways: differential expression analysis of RNA-Seq experiments using Rsubread and the edgeR quasi-likelihood pipeline. F1000Res. 2016 Jun 20;5:1438.

Cherry JJ, DiDonato CJ, Androphy EJ, Calo A, Potter K, Custer SK, Du S, Foley TL, Gopalsamy A, Reedich EJ, Gordo SM, Gordon W, Hosea N, Jones LH, Krizay DK, LaRosa G, Li H, Mathur S, Menard CA, Patel P, Ramos-Zayas R, Rietz A, Rong H, Zhang B, Tones MA. In vitro and in vivo effects of 2,4 diaminoquinazoline inhibitors of the decapping scavenger enzyme DcpS: Context-specific modulation of SMN transcript levels. PLoS One. 2017 Sep 25;12(9):e0185079.

Chiriboga CA, Swoboda KJ, Darras BT, Iannaccone ST, Montes J, De Vivo DC, Norris DA, Bennett CF, Bishop KM. Results from a phase 1 study of nusinersen (ISIS-SMN(Rx)) in children with spinal muscular atrophy. Neurology. 2016 Mar 8;86(10):890–7.

Chiriboga CA, Bruno C, Duong T, Fischer D, Mercuri E, Kirschner J, Kostera-Pruszczyk A, Jaber B, Gorni K, Kletzl H, Carruthers I, Martin C, Warren F, Scalco RS, Wagner KR, Muntoni F; JEWELFISH Study Group. Risdiplam in Patients Previously Treated with Other Therapies for Spinal Muscular Atrophy: An Interim Analysis from the JEWELFISH Study. Neurol Ther. 2023 Apr;12(2):543–557.

Corti S, Nizzardo M, Nardini M, Donadoni C, Salani S, Ronchi D, Saladino F, Bordoni A, Fortunato F, Del Bo R, Papadimitriou D, Locatelli F, Menozzi G, Strazzer S, Bresolin N, Comi GP. Neural stem cell transplantation can ameliorate the phenotype of a mouse model of spinal muscular atrophy. J Clin Invest. 2008 Oct;118(10):3316–30.

Darras BT, Crawford TO, Finkel RS, Mercuri E, De Vivo DC, Oskoui M, Tizzano EF, Ryan MM, Muntoni F, Zhao G, Staropoli J, McCampbell A, Petrillo M, Stebbins C, Fradette S, Farwell W, Sumner CJ. Neurofilament as a potential biomarker for spinal muscular atrophy. Ann Clin Transl Neurol. 2019 Apr 17;6(5):932–944.

Deguise MO, Baranello G, Mastella C, Beauvais A, Michaud J, Leone A, De Amicis R, Battezzati A, Dunham C, Selby K, Warman Chardon J, McMillan HJ, Huang YT, Courtney NL, Mole AJ, Kubinski S, Claus P, Murray LM, Bowerman M, Gillingwater TH, Bertoli S, Parson SH, Kothary R. Abnormal fatty acid metabolism is a core component of spinal muscular atrophy. Ann Clin Transl Neurol. 2019 Aug;6(8):1519–1532.

Deng C, Reinhard S, Hennlein L, Eilts J, Sachs S, Doose S, Jablonka S, Sauer M, Moradi M, Sendtner M. Impaired dynamic interaction of axonal endoplasmic reticulum and ribosomes contributes to defective stimulus-response in spinal muscular atrophy. Transl Neurodegener. 2022 Jun 2;11(1):31.

Detering NT, Zambon A, Hensel N, Kothary R, Swoboda K, Gillingwater TH, Baranello G; Workshop participants; Industry participants. 264th ENMC International Workshop: Multi-system involvement in spinal muscular atrophy Hoofddorp, the Netherlands, November 19th - 21st 2021. Neuromuscul Disord. 2022 Aug;32(8):697–705.

DiDonato CJ, Lorson CL, De Repentigny Y, Simard L, Chartrand C, Androphy EJ, Kothary R. Regulation of murine survival motor neuron (Smn) protein levels by modifying Smn exon 7 splicing. Hum Mol Genet. 2001 Nov 1;10(23):2727–36.

Doktor TK, Hua Y, Andersen HS, Brøner S, Liu YH, Wieckowska A, Dembic M, Bruun GH, Krainer AR, Andresen BS. RNA-sequencing of a mouse-model of spinal muscular atrophy reveals tissue-wide changes in splicing of U12-dependent introns. Nucleic Acids Res. 2017 Jan 9;45(1):395–416.

Eshraghi M, McFall E, Gibeault S, Kothary R. Effect of genetic background on the phenotype of the Smn2B/- mouse model of spinal muscular atrophy. Hum Mol Genet. 2016 Oct 15;25(20):4494–4506.

Errico F, Marino C, Grimaldi M, Nuzzo T, Bassareo V, Valsecchi V, Panicucci C, Di Schiavi E, Mazza T, Bruno C, D’Amico A, Carta M, D’Ursi AM, Bertini E, Pellizzoni L, Usiello A. Nusinersen Induces Disease-Severity-Specific Neurometabolic Effects in Spinal Muscular Atrophy. Biomolecules. 2022 Oct 6;12(10):1431.

Fajardo-Jiménez MJ, Tejada-Moreno JA, Mejía-García A, Villegas-Lanau A, Zapata-Builes W, Restrepo JE, Cuartas GP, Hernandez JC. Ehlers-Danlos: A Literature Review and Case Report in a Colombian Woman with Multiple Comorbidities. Genes (Basel). 2022 Nov 15;13(11):2118.

Fallini C, Donlin-Asp PG, Rouanet JP, Bassell GJ, Rossoll W. Deficiency of the Survival of Motor Neuron Protein Impairs mRNA Localization and Local Translation in the Growth Cone of Motor Neurons. J Neurosci. 2016 Mar 30;36(13):3811–20.

Faravelli I, Meneri M, Saccomanno D, Velardo D, Abati E, Gagliardi D, Parente V, Petrozzi L, Ronchi D, Stocchetti N, Calderini E, D’Angelo G, Chidini G, Prandi E, Ricci G, Siciliano G, Bresolin N, Comi GP, Corti S, Magri F, Govoni A. Nusinersen treatment and cerebrospinal fluid neurofilaments: An explorative study on Spinal Muscular Atrophy type 3 patients. J Cell Mol Med. 2020 Mar;24(5):3034–3039.

Flotats-Bastardas M, Bitzan L, Grell C, Martakis K, Winter B, Zemlin M, Wurster CD, Uzelac Z, Weiß C, Hahn A. Paradoxical increase of neurofilaments in SMA patients treated with onasemnogene abeparvovec-xioi. Front Neurol. 2023 Dec 13;14:1269406.

Finkel RS, Mercuri E, Darras BT, Connolly AM, Kuntz NL, Kirschner J, Chiriboga CA, Saito K, Servais L, Tizzano E, Topaloglu H, Tulinius M, Montes J, Glanzman AM, Bishop K, Zhong ZJ, Gheuens S, Bennett CF, Schneider E, Farwell W, De Vivo DC; ENDEAR Study Group. Nusinersen versus Sham Control in Infantile-Onset Spinal Muscular Atrophy. N Engl J Med. 2017 Nov 2;377(18):1723–1732.

Glascock J, Darras BT, Crawford TO, Sumner CJ, Kolb SJ, DiDonato C, Elsheikh B, Howell K, Farwell W, Valente M, Petrillo M, Tingey J, Jarecki J. Identifying Biomarkers of Spinal Muscular Atrophy for Further Development. J Neuromuscul Dis. 2023;10(5):937–954.

Groen EJN, Talbot K, Gillingwater TH. Advances in therapy for spinal muscular atrophy: promises and challenges. Nat Rev Neurol. 2018 Apr;14(4):214–224. doi: 10.1038/nrneurol.2018.4.

Hammond SM, Gogliotti RG, Rao V, Beauvais A, Kothary R, DiDonato CJ. Mouse survival motor neuron alleles that mimic SMN2 splicing and are inducible rescue embryonic lethality early in development but not late. PLoS One. 2010 Dec 29;5(12):e15887.

Hensel N, Claus P. The Actin Cytoskeleton in SMA and ALS: How Does It Contribute to Motoneuron Degeneration? Neuroscientist. 2018 Feb;24(1):54–72.

Hensel N, Brickwedde H, Tsaknakis K, Grages A, Braunschweig L, Lüders KA, Lorenz HM, Lippross S, Walter LM, Tavassol F, Lienenklaus S, Neunaber C, Claus P, Hell AK. Altered bone development with impaired cartilage formation precedes neuromuscular symptoms in spinal muscular atrophy. Hum Mol Genet. 2020 Sep 29;29(16):2662–2673.

Iannarone VJ, Cruz GE, Hilliard BA, Barbe MF. The answer depends on the question: Optimal conditions for western blot characterization of muscle collagen type 1 depends on desired isoform. J Biol Methods. 2019 Aug 15;6(3):e117.

James R, Chaytow H, Ledahawsky LM, Gillingwater TH. Revisiting the role of mitochondria in spinal muscular atrophy. Cell Mol Life Sci. 2021 May;78(10):4785–4804.

Kessler T, Latzer P, Schmid D, Warnken U, Saffari A, Ziegler A, Kollmer J, Möhlenbruch M, Ulfert C, Herweh C, Wildemann B, Wick W, Weiler M. Cerebrospinal fluid proteomic profiling in nusinersen-treated patients with spinal muscular atrophy. J Neurochem. 2020 Jun;153(5):650–661.

Khatri IA, Chaudhry US, Seikaly MG, Browne RH, Iannaccone ST. Low bone mineral density in spinal muscular atrophy. J Clin Neuromuscul Dis. 2008 Sep;10(1):11–7.

Kolb SJ, Coffey CS, Yankey JW, Krosschell K, Arnold WD, Rutkove SB, Swoboda KJ, Reyna SP, Sakonju A, Darras BT, Shell R, Kuntz N, Castro D, Parsons J, Connolly AM, Chiriboga CA, McDonald C, Burnette WB, Werner K, Thangarajh M, Shieh PB, Finanger E, Cudkowicz ME, McGovern MM, McNeil DE, Finkel R, Iannaccone ST, Kaye E, Kingsley A, Renusch SR, McGovern VL, Wang X, Zaworski PG, Prior TW, Burghes AHM, Bartlett A, Kissel JT; NeuroNEXT Clinical Trial Network on behalf of the NN101 SMA Biomarker Investigators. Natural history of infantile-onset spinal muscular atrophy. Ann Neurol. 2017 Dec;82(6):883–891.

Lauria F, Bernabò P, Tebaldi T, Groen EJN, Perenthaler E, Maniscalco F, Rossi A, Donzel D, Clamer M, Marchioretto M, Omersa N, Orri J, Dalla Serra M, Anderluh G, Quattrone A, Inga A, Gillingwater TH, Viero G. SMN-primed ribosomes modulate the translation of transcripts related to spinal muscular atrophy. Nat Cell Biol. 2020 Oct;22(10):1239–1251.

Lauria F, Tebaldi T, Bernabò P, Groen EJN, Gillingwater TH, Viero G. riboWaltz: Optimization of ribosome P-site positioning in ribosome profiling data.

Lefebvre S, Bürglen L, Reboullet S, Clermont O, Burlet P, Viollet L, Benichou B, Cruaud C, Millasseau P, Zeviani M, et al. Identification and characterization of a spinal muscular atrophy-determining gene. Cell. 1995 Jan 13;80(1):155–65.

Lefebvre S, Burlet P, Liu Q, Bertrandy S, Clermont O, Munnich A, Dreyfuss G, Melki J. Correlation between severity and SMN protein level in spinal muscular atrophy. Nat Genet. 1997 Jul;16(3):265–9.

Leow DM, Ng YK, Wang LC, Koh HW, Zhao T, Khong ZJ, Tabaglio T, Narayanan G, Giadone RM, Sobota RM, Ng SY, Teo AKK, Parson SH, Rubin LL, Ong WY, Darras BT, Yeo CJ. Hepatocyte-intrinsic SMN deficiency drives metabolic dysfunction and liver steatosis in spinal muscular atrophy. J Clin Invest. 2024 May 9;134(12):e173702.

Lorson CL, Hahnen E, Androphy EJ, Wirth B. A single nucleotide in the SMN gene regulates splicing and is responsible for spinal muscular atrophy. Proc Natl Acad Sci U S A. 1999 May 25;96(11):6307–11.

Lotti F, Imlach WL, Saieva L, Beck ES, Hao le T, Li DK, Jiao W, Mentis GZ, Beattie CE, McCabe BD, Pellizzoni L. An SMN-dependent U12 splicing event essential for motor circuit function. Cell. 2012 Oct 12;151(2):440–54.

Lun AT, Chen Y, Smyth GK. It’s DE-licious: A Recipe for Differential Expression Analyses of RNA-seq Experiments Using Quasi-Likelihood Methods in edgeR. Methods Mol Biol. 2016;1418:391–416.

Maretina M, Koroleva V, Shchugareva L, Glotov A, Kiselev A. The Relevance of Spinal Muscular Atrophy Biomarkers in the Treatment Era. Biomedicines. 2024 Oct 30;12(11):2486. doi: 10.3390/biomedicines12112486.

McCarthy DJ, Chen Y, Smyth GK. Differential expression analysis of multifactor RNA-Seq experiments with respect to biological variation. Nucleic Acids Res. 2012 May;40(10):4288–97.

Mendell JR, Al-Zaidy S, Shell R, Arnold WD, Rodino-Klapac LR, Prior TW, Lowes L, Alfano L, Berry K, Church K, Kissel JT, Nagendran S, L’Italien J, Sproule DM, Wells C, Cardenas JA, Heitzer MD, Kaspar A, Corcoran S, Braun L, Likhite S, Miranda C, Meyer K, Foust KD, Burghes AHM, Kaspar BK. Single-Dose Gene-Replacement Therapy for Spinal Muscular Atrophy. N Engl J Med. 2017 Nov 2;377(18):1713–1722.

Meneri M, Abati E, Gagliardi D, Faravelli I, Parente V, Ratti A, Verde F, Ticozzi N, Comi GP, Ottoboni L, Corti S. Identification of Novel Biomarkers of Spinal Muscular Atrophy and Therapeutic Response by Proteomic and Metabolomic Profiling of Human Biological Fluid Samples. Biomedicines. 2023 Apr 23;11(5):1254.

Mercuri E, Cicala G, Villa M, Pera MC. What did we learn from new treatments in SMA? A narrative review. Acta Myol. 2025 Mar;44(1):28–32.

Monani UR, Lorson CL, Parsons DW, Prior TW, Androphy EJ, Burghes AH, McPherson JD. A single nucleotide difference that alters splicing patterns distinguishes the SMA gene SMN1 from the copy gene SMN2. Hum Mol Genet. 1999 Jul;8(7):1177–83.

Motyl AAL, Faller KME, Groen EJN, Kline RA, Eaton SL, Ledahawsky LM, Chaytow H, Lamont DJ, Wishart TM, Huang YT, Gillingwater TH. Pre-natal manifestation of systemic developmental abnormalities in spinal muscular atrophy. Hum Mol Genet. 2020 Sep 29;29(16):2674–2683. doi: 10.1093/hmg/ddaa146

Ngawa M, Dal Farra F, Marinescu AD, Servais L. Longitudinal developmental profile of newborns and toddlers treated for spinal muscular atrophy. Ther Adv Neurol Disord. 2023 Feb 20;16:17562864231154335.

Nichterwitz S, Nijssen J, Storvall H, Schweingruber C, Comley LH, Allodi I, Lee MV, Deng Q, Sandberg R, Hedlund E. LCM-seq reveals unique transcriptional adaptation mechanisms of resistant neurons and identifies protective pathways in spinal muscular atrophy. Genome Res. 2020 Aug;30(8):1083–1096. doi: 10.1101/gr.265017.120.

Nishio H, Niba ETE, Saito T, Okamoto K, Takeshima Y, Awano H. Spinal Muscular Atrophy: The Past, Present, and Future of Diagnosis and Treatment. Int J Mol Sci. 2023 Jul 26;24(15):11939.

Olaso R, Joshi V, Fernandez J, Roblot N, Courageot S, Bonnefont JP, Melki J. Activation of RNA metabolism-related genes in mouse but not human tissues deficient in SMN. Physiol Genomics. 2006 Jan 12;24(2):97–104.

Otsuki N, Arakawa R, Kaneko K, Aoki R, Arakawa M, Saito K. A new biomarker candidate for spinal muscular atrophy: Identification of a peripheral blood cell population capable of monitoring the level of survival motor neuron protein. PLoS One. 2018 Aug 13;13(8):e0201764.

Poirier A, Weetall M, Heinig K, Bucheli F, Schoenlein K, Alsenz J, Bassett S, Ullah M, Senn C, Ratni H, Naryshkin N, Paushkin S, Mueller L. Risdiplam distributes and increases SMN protein in both the central nervous system and peripheral organs. Pharmacol Res Perspect. 2018 Nov 29;6(6):e00447. doi: 10.1002/prp2.447.

Price PL, Morderer D, Rossoll W. RNP Assembly Defects in Spinal Muscular Atrophy. Adv Neurobiol. 2018;20:143–171

Ramos DM, d’Ydewalle C, Gabbeta V, Dakka A, Klein SK, Norris DA, Matson J, Taylor SJ, Zaworski PG, Prior TW, Snyder PJ, Valdivia D, Hatem CL, Waters I, Gupte N, Swoboda KJ, Rigo F, Bennett CF, Naryshkin N, Paushkin S, Crawford TO, Sumner CJ. Age-dependent SMN expression in disease-relevant tissue and implications for SMA treatment. J Clin Invest. 2019 Nov 1;129(11):4817–4831.

Reedich EJ, Kalski M, Armijo N, Cox GA, DiDonato CJ. Spinal motor neuron loss occurs through a p53- and-p21-independent mechanism in the Smn2B/- mouse model of spinal muscular atrophy. Exp Neurol. 2021 Mar;337:113587.

Reilly A, Beauvais A, Al-Aarg M, Yaworski R, Sutton ER, Thebault S, Kothary R. Peripheral defects precede neuromuscular pathology in the Smn2B/- mouse model of spinal muscular atrophy. J Neuromuscul Dis. 2024 Nov;11(6):1200–1210.

Rossoll W, Jablonka S, Andreassi C, Kröning AK, Karle K, Monani UR, Sendtner M. Smn, the spinal muscular atrophy-determining gene product, modulates axon growth and localization of beta-actin mRNA in growth cones of motoneurons. J Cell Biol. 2003 Nov 24;163(4):801–12.

Ruggiu M, McGovern VL, Lotti F, Saieva L, Li DK, Kariya S, Monani UR, Burghes AH, Pellizzoni L. A role for SMN exon 7 splicing in the selective vulnerability of motor neurons in spinal muscular atrophy. Mol Cell Biol. 2012 Jan;32(1):126–38.

Sanchez G, Dury AY, Murray LM, Biondi O, Tadesse H, El Fatimy R, Kothary R, Charbonnier F, Khandjian EW, Côté J. A novel function for the survival motoneuron protein as a translational regulator. Hum Mol Genet. 2013 Feb 15;22(4):668–84.

Sharma G, Paganin M, Lauria F, Perenthaler E, Viero G. The SMN-ribosome interplay: a new opportunity for Spinal Muscular Atrophy therapies. Biochem Soc Trans. 2024 Feb 28;52(1):465–479.

Signoria I, Zwartkruis MM, Geerlofs L, Perenthaler E, Faller KME, James R, McHale-Owen H, Green JW, Kortooms J, Snellen SH, Asselman FL, Gillingwater TH, Viero G, Wadman RI, van der Pol WL, Groen EJN. Patient-specific responses to SMN2 splice-modifying treatments in spinal muscular atrophy fibroblasts. Mol Ther Methods Clin Dev. 2024 Nov 13;32(4):101379.

Somers E, Lees RD, Hoban K, Sleigh JN, Zhou H, Muntoni F, Talbot K, Gillingwater TH, Parson SH. Vascular Defects and Spinal Cord Hypoxia in Spinal Muscular Atrophy. Ann Neurol. 2016 Feb;79(2):217–30. doi: 10.1002/ana.24549.

Sonenberg N, Hinnebusch AG. Regulation of translation initiation in eukaryotes: mechanisms and biological targets. Cell. 2009 Feb 20;136(4):731–45.

Susanto TT, Hung V, Levine AG, Chen Y, Kerr CH, Yoo Y, Oses-Prieto JA, Fromm L, Zhang Z, Lantz TC, Fujii K, Wernig M, Burlingame AL, Ruggero D, Barna M. RAPIDASH: Tag-free enrichment of ribosome-associated proteins reveals composition dynamics in embryonic tissue, cancer cells, and macrophages. Mol Cell. 2024 Sep 19;84(18):3545–3563.e25.

Tadesse H, Deschênes-Furry J, Boisvenue S, Côté J. KH-type splicing regulatory protein interacts with survival motor neuron protein and is misregulated in spinal muscular atrophy. Hum Mol Genet. 2008 Feb 15;17(4):506–24.

Tapken I, Schweitzer T, Paganin M, Schüning T, Detering NT, Sharma G, Niesert M, Saffari A, Kuhn D, Glynn A, Cieri F, Santonicola P, Cannet C, Gerstner F, Faller KME, Huang YT, Kothary R, Gillingwater TH, Di Schiavi E, Simon CM, Hensel N, Ziegler A, Viero G, Pich A, Claus P. The systemic complexity of a monogenic disease: the molecular network of spinal muscular atrophy. Brain. 2025 Feb 3;148(2):580–596.

Tosi M, Cumbo F, Catteruccia M, Carlesi A, Mizzoni I, De Luca G, Cherchi C, Cutrera R, Bertini E, D’Amico A. Neurocognitive profile of a cohort of SMA type 1 pediatric patients and emotional aspects, resilience and coping strategies of their caregivers. Eur J Paediatr Neurol. 2023 Mar;43:36–43.

Uhlén M, Fagerberg L, Hallström BM, Lindskog C, Oksvold P, Mardinoglu A, Sivertsson Å, Kampf C, Sjöstedt E, Asplund A, Olsson I, Edlund K, Lundberg E, Navani S, Szigyarto CA, Odeberg J, Djureinovic D, Takanen JO, Hober S, Alm T, Edqvist PH, Berling H, Tegel H, Mulder J, Rockberg J, Nilsson P, Schwenk JM, Hamsten M, von Feilitzen K, Forsberg M, Persson L, Johansson F, Zwahlen M, von Heijne G, Nielsen J, Pontén F. Proteomics. Tissue-based map of the human proteome. Science. 2015 Jan 23;347(6220):1260419.

Vai S, Bianchi ML, Moroni I, Mastella C, Broggi F, Morandi L, Arnoldi MT, Bussolino C, Baranello G. Bone and Spinal Muscular Atrophy. Bone. 2015 Oct;79:116-20. doi: 10.1016/j.bone.2015.05.039.

Vitte JM, Davoult B, Roblot N, Mayer M, Joshi V, Courageot S, Tronche F, Vadrot J, Moreau MH, Kemeny F, Melki J. Deletion of murine Smn exon 7 directed to liver leads to severe defect of liver development associated with iron overload. Am J Pathol. 2004 Nov;165(5):1731–41.

Wadman RI, Stam M, Jansen MD, van der Weegen Y, Wijngaarde CA, Harschnitz O, Sodaar P, Braun KP, Dooijes D, Lemmink HH, van den Berg LH, van der Pol WL. A Comparative Study of SMN Protein and mRNA in Blood and Fibroblasts in Patients with Spinal Muscular Atrophy and Healthy Controls. PLoS One. 2016 Nov 28;11(11):e0167087.

Wadman RI, Jansen MD, Stam M, Wijngaarde CA, Curial CAD, Medic J, Sodaar P, Schouten J, Vijzelaar R, Lemmink HH, van den Berg LH, Groen EJN, van der Pol WL. Intragenic and structural variation in the SMN locus and clinical variability in spinal muscular atrophy. Brain Commun. 2020 Jun 8;2(2):fcaa075.

Wasserman HM, Hornung LN, Stenger PJ, Rutter MM, Wong BL, Rybalsky I, Khoury JC, Kalkwarf HJ. Low bone mineral density and fractures are highly prevalent in pediatric patients with spinal muscular atrophy regardless of disease severity. Neuromuscul Disord. 2017 Apr;27(4):331–337.

Wirth B. Spinal Muscular Atrophy: In the Challenge Lies a Solution. Trends Neurosci. 2021 Apr;44(4):306–322.

Yeo CJJ, Tizzano EF, Darras BT. Challenges and opportunities in spinal muscular atrophy therapeutics. Lancet Neurol. 2024 Feb;23(2):205–218.

Yeo CJJ, Darras BT. Overturning the Paradigm of Spinal Muscular Atrophy as Just a Motor Neuron Disease. Pediatr Neurol. 2020 Aug;109:12–19

Zhang Z, Lotti F, Dittmar K, Younis I, Wan L, Kasim M, Dreyfuss G. SMN deficiency causes tissue-specific perturbations in the repertoire of snRNAs and widespread defects in splicing. Cell. 2008 May 16;133(4):585–600.

Zhou H, Hong Y, Scoto M, Thomson A, Pead E, MacGillivray T, Hernandez-Gerez E, Catapano F, Meng J, Zhang Q, Hunter G, Shorrock HK, Ng TK, Hamida A, Sanson M, Baranello G, Howell K, Gillingwater TH, Brogan P, Thompson DA, Parson SH, Muntoni F. Microvasculopathy in spinal muscular atrophy is driven by a reversible autonomous endothelial cell defect. J Clin Invest. 2022 Nov 1;132(21):e153430.

